# Chromatin compaction upon CDK12 inhibition drives long gene silencing and a combinatorial lethal dependency on PAF1

**DOI:** 10.64898/2026.01.27.701921

**Authors:** Jing Liang, Harri M Itkonen

**Affiliations:** Department of Biochemistry and Developmental Biology, Faculty of Medicine, University of Helsinki, 00014 Helsinki, Finland; Department of Clinical Molecular Biology, EpiGen, Institute of Clinical Medicine, University of Oslo, 1171 Oslo, Norway; EpiGen, Medical Division, Akershus University Hospital, 1478 Lørenskog, Norway

**Keywords:** Transcriptional kinases, chromatin accessibility, cyclin-dependent kinase 12/13, transcriptionally active RNA Pol II complex

## Abstract

Loss-of-function mutations in cyclin-dependent kinase 12 (CDK12) define an aggressive subtype of metastatic castration-resistant prostate cancer (mCRPC) characterized by genomic instability, and depletion of CDK12 activity downregulates the expression of long DNA repair genes. While these transcriptional defects are well-documented, potential alterations in chromatin that may topologically affect this silencing and the subsequent adaptive survival mechanisms of tumor cells remain poorly understood. Here, we employ a dynamic multi-omics strategy to map the remodeling of the RNA Polymerase II (RNA Pol II) complex and chromatin landscape. We utilized 5-ethynyl uridine (5-EU) metabolic labeling combined with mass spectrometry (MS) to specifically capture the transcriptionally active RNA Pol II complex, distinguishing active elongation complexes from static background, and map how its interactome changes when the activity of the major transcriptional kinases CDK7, CDK9 or CDK12/13 is inhibited. These data are then integrated with ATAC-seq, transcriptomics, and targeted synthetic lethality screen to validate findings from the RNA Pol II capture. We show that, unlike CDK9 inhibition, CDK12/13 inhibition triggers a progressive chromatin compaction. This is characterized by the decrease in the binding activity of CTCF to chromosomes closing, which acts as a physical blockade specifically silencing long genes when CDK12 activity is lost. In response to this topological remodeling, cells launch an adaptive survival program by recruiting the transcription elongation factor PAF1 and the DNA damage sensor DDB1 to the stalled RNA Pol II. We identify PAF1 as a critical vulnerability in this context. Collectively, we identify CDK12 as a regulator of chromatin topology rather than merely an elongation kinase. The chromatin compaction upon CDK12/13 inhibition creates a unique dependency on the PAF1-mediated adaptive remodeling. Specifically, targeting PAF1 converts this stress response into a lethal trap, establishing a potent, kinase-selective combination lethal strategy for *CDK12*-deficient prostate cancer.

## Introduction

Dysregulated transcriptional control is a fundamental hallmark of cancer, enabling malignant cells to tolerate oncogenic stress and sustain unrestricted proliferation [1, 2]. This abnormal gene expression program is highly dependent on the dynamic post-translational modifications of the RNA Polymerase II (RNA Pol II) C-terminal domain (CTD), which serves as a scaffold for recruiting RNA processing factors [3, 4]. The phosphorylation cycle of the CTD is strictly governed by the cyclin-dependent kinase (CDK) family, specifically the transcriptional CDKs [5–7]. Among these, CDK7 initiates transcription by phosphorylating Serine-5 (Ser5), while CDK9 facilitates the release of RNA Pol II from promoter-proximal pausing by phosphorylating Serine-2 (Ser2) and negative regulators of transcription [8–10]. Distinct from these, cyclin-dependent kinase 12 (CDK12) and CDK13, has emerged as a critical regulator specifically dedicated to productive transcriptional elongation [11, 12].

Prostate cancer (PC) is most common cancer in men, and *CDK12* mutations are potent driver of prostate cancer aggressiveness and progression [13, 14]. Loss-of-function mutations in *CDK12* are enriched in aggressive prostate cancer, especially metastatic castration-resistant prostate cancer (mCRPC) and identify a distinct subclass of tumors associated with rapid disease progression and poor prognosis [14]. Tumors harboring *CDK12* inactivation exhibit a distinct tandem duplicator phenotype (TDP), characterized by genome-wide focal tandem duplications [14, 15]. Mechanistically, *CDK12* loss compromises genomic stability, and its full activity is important for the expression of long genes, particularly those involved in the DNA damage response (DDR) such as *BRCA1* (breast cancer type 1) and *ATM* (ataxia-telangiectasia mutated) [11, 12, 16]. Downregulation of these long genes is proposed to be driven by premature intronic polyadenylation (IPA) due to the failure of RNA Pol II to suppress proximal cleavage sites [16, 17]. Although the genomic scars (TDP) and transcriptomic consequences have been well documented, the molecular sequence of events between CDK12 inactivation and the downregulation of long genes remains incompletely understood. It is hypothesized that CDK12 does not merely catalyze phosphorylation but structurally scaffolds the elongation machinery to suppress proximal splicing sites and maintain processivity [16, 18]. However, transcriptional output is not solely determined by the enzymatic activity of polymerase but is intrinsically coupled to the physical state of the chromatin template [19, 20]. These data therefore form an intriguing hypothesis that *CDK12* inactivation may also impose a global physical constraint on the chromatin template to contribute to the aggressive phenotype of cancer cells. Recent single-cell epigenomic profiling in PC has demonstrated that chromatin accessibility is a primary determinant of transcriptional reprogramming during drug resistance, preceding changes in gene expression [21]. Therefore, it is crucial to understand how transcription kinase inactivation affects chromatin structure and transcription machinery to affect transcription.

Elucidating these dynamic mechanisms is challenging because traditional antibody-based immunoprecipitation (IP) captures the total pool of RNA Pol II, which is dominated by static or stalled complexes [22]. This limitation obscures the rapid and active remodeling events that occur upon acute kinase inhibition [22, 23]. To overcome this, strategies that capture the nascent RNA interactome, such as capture of the newly transcribed RNA interactome (RICK) or metabolic labeling with 5-ethynyl uridine (5-EU), have been developed to specifically isolate proteins coupled to active transcription [22, 24]. These methods act as a “molecular sieve”, filtering out inactive background noise to specifically enrich protein complexes associated with nascent RNA and active transcription [22, 24, 25].

In this study, we aimed to systematically map the dynamic remodeling of the RNA Pol II complex and chromatin structure in PC cells following inhibition of major transcriptional kinases using a systematic multi-omics strategy. We reveal that inhibition of major transcriptional kinases induces distinct alterations in chromatin accessibility. In contrast to CDK7 and CDK9 inhibition, CDK12 inhibition triggers widespread “chromatin condensation” program. This chromatin compaction constitutes a structural blockade that specifically silences long genes, including those involved in DNA damage responses (DDR). Similarly, this compression-induced downregulation of long genes also observed in PC patient samples with *CDK12* truncating mutations. Furthermore, our dynamic interactome data, captured via 5-EU metabolic labeling, uncovered an adaptive survival response. CDK12-deficient cells recruit DNA repair and remodeling factors, centered on PAF1 and DDB1, to the RNA Pol II machinery to adapt to the stress. Finally, we demonstrate that targeting this PAF1-mediated adaptive pathway induces combinatorial lethality, thereby providing a novel mechanistic rationale and specific therapeutic strategy exploiting synthetic lethality against aggressive *CDK12*-mutant PC tumors.

## Results

### 5-EU metabolic labelling of RNA captures transcriptionally-engaged RNA Pol II complex

To investigate the real-time impact of inhibiting major transcriptional kinases on the dynamics of transcription complexes, it is essential to distinguish “elongating active complexes” from “static complexes”. Traditional antibody-based immunoprecipitation (IP) captures the total pool of RNA Pol II and its stably associated partners, making it difficult to precisely reflect the dynamic remodeling that occurs following kinase inhibition. To address this, we adapted the principles of the “RICK” technique, originally designed for capturing the nascent RNA interactome and Isolation of Proteins on Nascent DNA (iPOND) principle [22, 25, 26], to establish an optimized strategy. In order to enrich RNA Pol II rather than other protein machineries assembled on the mRNA later during splicing, transport, and translation, the labeling time should be short. Therefore, we used a 10-minute pulse of 5-ethynyl uridine (5-EU) to label nascent RNA, followed by a biotin-streptavidin system to specifically isolate protein complexes directly coupled to nascent transcripts (**Fig. 1A**). To confirm this method, we first evaluated whether 5-EU metabolic labeling could selectively enrich for transcriptionally active RNA Pol II. As we expected, streptavidin affinity purification following a 10-minute 5-EU pulse resulted in enrichment of RNA Pol II. In contrast, negative control GAPDH showed no enrichment, confirming high specificity (**Fig. 1B**).

**Figure 1.**
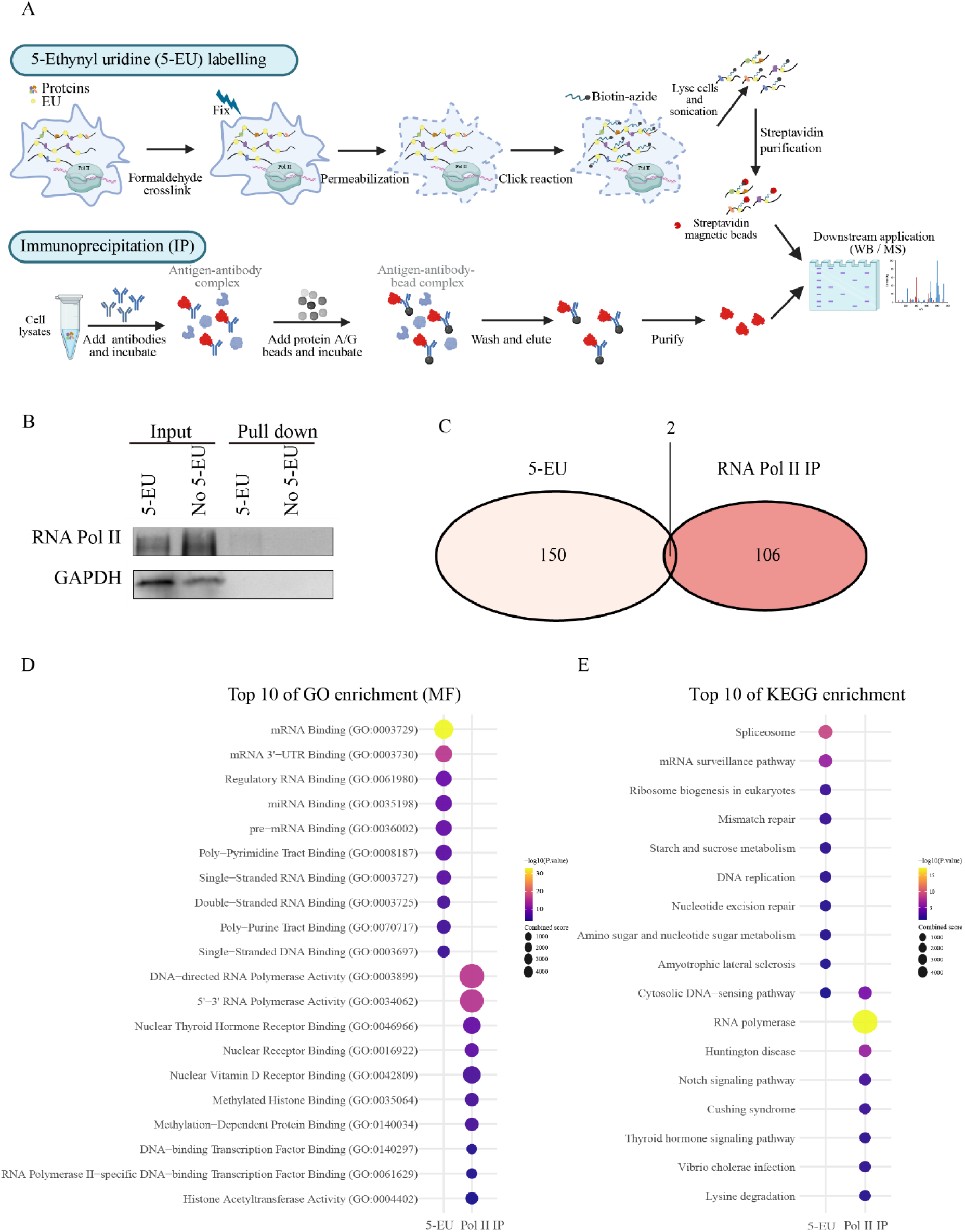
5-EU metabolic labeling specifically captures the transcriptionally active RNA Pol II interactome. **A.** Schematic representation of the 5-EU metabolic labeling strategy. Cells were treated with 1 mM 5-ethynyl uridine (5-EU) for 10 min to label nascent RNA, followed by biotinylation (biotin-azide) via click chemistry and streptavidin-based affinity purification to enrich proteins. **B** Western blot validation of the 5-EU enrichment strategy. C4-2 cells were labeled with 5-EU for 10 minutes and then enriched according to the procedure in this study. The final eluent was analyzed by Western blotting using antibodies against RNA Pol II and GAPDH. Non-labeled cells served as negative control. **C** Venn diagram showing the overlap and specificity of proteins identified by 5-EU enrichment versus RNA Pol II immunoprecipitation (IP). **D** Gene Ontology (GO) and **E** KEGG enrichment analysis of proteins unique to the 5-EU and RNA Pol II IP datasets using enrichr [72].

To directly compare the “elongating active complexes” and the “static complexes”, we performed mass spectrometry (MS) analysis on samples enriched via both 5-EU labeling and traditional IP strategies. Comparative analysis identified 152 proteins significant enriched by 5-EU labeling and 108 significant proteins enriched by RNA Pol II IP (**Supplementary Table 1**). To further elucidate the distinctions between these two methods, we performed an intersection analysis of the datasets. Consistent with our hypothesis, a direct comparison revealed minimal overlap, with only two proteins shared between the groups; 150 proteins were unique to the 5-EU enrichment, while 106 were unique to the RNA Pol II IP (**Fig. 1C**). To further characterize the functional profiles captured by each strategy, we conducted Gene Ontology (GO) and Kyoto Encyclopedia of Genes and Genomes (KEGG) enrichment analyses on the unique proteins identified by each method. GO analysis revealed that the 5-EU fraction was specifically enriched for dynamic factors tightly coupled to transcriptional elongation, including RNA splicing (GO:0003729), mRNA binding (GO:0003729), and pre-mRNA binding (GO:0036002) (**Fig. 1D**). These findings show that labelling of the nascent transcription captures processes involved in the initial mRNA maturation rather than later steps, such as the ribosome. Similarly, KEGG analysis showed significant enrichment of the spliceosome and mRNA surveillance pathways (**Fig. 1E**), which regulate pre-mRNA splicing and quality control, and are key links in co- and post-transcriptional mRNA processing [27].. Collectively, these results indicate that the 5-EU-enriched proteome is biased towards factors mediating nascent RNA maturation, stability, and translational regulation. Conversely, the RNA Pol II IP dataset showed greater enrichment for static DNA-binding factors (**Fig. 1D**), such as catalytic activity (GO:0003899) and DNA binding (GO:0034062), reflecting the basal catalytic machinery of RNA Pol II [28]. The enrichment of terms such as Nuclear Receptor Binding (GO:0016922), DNA-binding Transcription Factor Binding (GO:0140297), and RNA Pol II-specific DNA-binding Transcription Factor Binding (GO:0061629) (**Fig. 1D**) further suggests that the Pol II immunoprecipitated predominantly captures transcription factors and co-regulators recruited to the polymerase and / or chromatin during transcription initiation, which is known to be slow process relative to transcription elongation [29].

These findings establish 5-EU based labeling of transcriptionally engaged RNA Pol II as a high-resolution “molecular sieve” capable of filtering background noise to specifically isolate protein complexes associated with nascent RNA and active transcription. Moreover, we provide the first direct proteomic comparison between 5-EU-labeled RNA affinity purification and RNA Pol II immunoprecipitation. This combined strategy offers a robust platform for dissecting how inhibition of the major transcriptional kinases affect the composition of the transcriptional machinery associated with active polymerases.

### CDK12 inhibition induces chromatin condensation signature and downregulates long genes

To understand how inhibition of transcriptional regulatory kinases reshapes the RNA Pol II transcriptional landscape, we first evaluated changes at the RNA level. We found that treatment with different transcription kinase inhibitors resulted in their respective unique transcriptomics programs (**Fig. 2A**). Subsequently, we identified differentially expressed genes (DEGs) across treatment groups (**Suppl. Figure 1**) and performed functional enrichment analysis. The results revealed that after 4 hours of treatment, the CDK12 inhibition group was specifically enriched for biological pathways related to “chromosome condensation” (**Fig. 2B**). In contrast, CDK7 or CDK9 inhibition primarily resulted in the downregulation of the holoenzyme complex or downregulation of pathways related to transcription (**Suppl. Figure 2**). Specifically, CDK9 inhibition (NVP-2) led directly to the downregulation of transcriptional regulation pathways, consistent with the loss of its function as a positive transcriptional elongation factor (**Suppl. Figure 2A**). Meanwhile, CDK7 inhibition (YKL-5-124) mainly caused the downregulation of genes associated with the “holoenzyme complex” (**Suppl. Figure 2B**), aligning with its role as a transcription initiation kinase and reflecting a collapse of the core transcriptional machinery [30]. Our previous research showed that CDK12/13 inhibitor (THZ531) treatment for 4 hours resulted in a decrease in the acetylation of histone 3 [31] and similar effects on the same histone mark has also been reported by others [32]. These findings align with the early “condensation” instruction observed in our enrichment analysis following *CDK12* inhibition in this study. This early induction of chromatin condensation may come with functional consequences.

**Figure 2.**
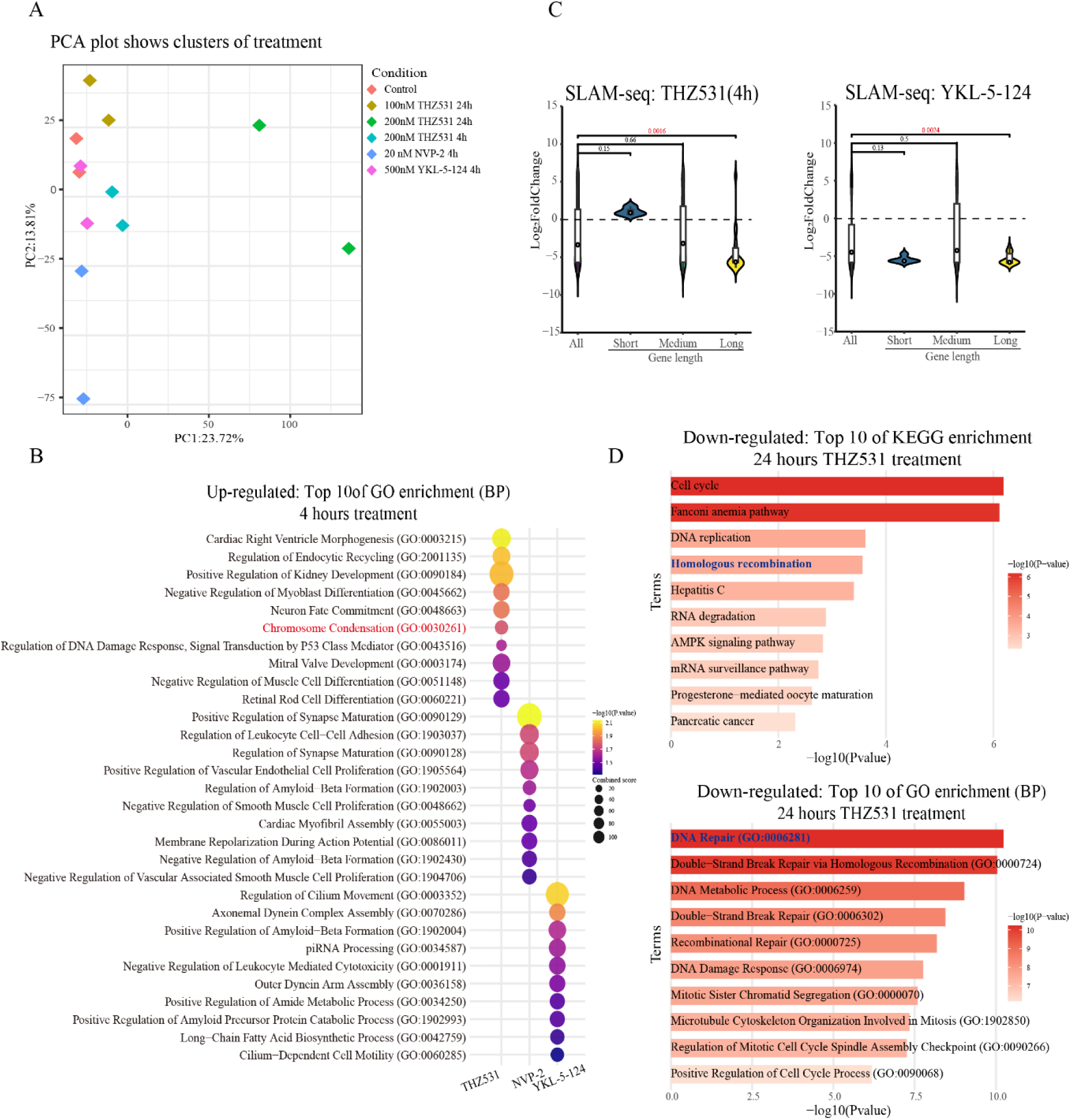
CDK12/13 inhibition induces chromosome condensation transcriptional program and specific downregulation of long genes. **A** Principal-component analysis (PCA) of the transcriptional profiling. The PCA figure represents a two-dimensional scatterplot of the first two principal components of the SLAM-seq data. Different inhibitor treatment groups at different time points are represented by different colors as indicated by the legend provided within the graph. Each dot represents a biological replicate of a sample. **B** GO enrichment analysis of genes downregulated upon CDK7 (YKL-5-124, 500 nM), CDK9 (NVP-2, 20 nM), and CDK12 (THZ531, 200 nM) inhibition for 4 hours treatment. The chromosome condensation entry is significantly enriched in biological process (BP) in CDK12 inhibition, while those in CDK7 and CDK9 were not. **C** CDK12 inhibition significantly downregulated the expression of long genes after 4 and 24 hours (SLAM-DUNK). Gene length stratification was consistent with our previous studies [18], and significance was assessed using Student’s *t*-test. **D** CDK12/13 inhibition leads to a significant enrichment of DNA repair (GO BP) and homologous recombination (KEGG) in downregulated genes.

*CDK12* inhibition leads to downregulation of transcription. We and others have shown that CDK12 inhibition stimulates transcription of short genes and downregulates of long genes [18, 31]. Here, our gene length-dependent analysis showed that CDK12 inhibition caused specific downregulation of long genes, consistent with the previous findings. Notably, this defect was progressively exacerbated as the treatment duration extended from 4 to 24 hours (**Fig. 2C**). For comparison, we subjected the CDK7 and CDK9 inhibition groups to the same analysis. Our data revealed that CDK7 inhibition, in addition to causing global downregulation, also significantly downregulated the expression of long genes (**Suppl. Figure 3A**). Considering the functional enrichment results (**Suppl. Figure 2B**), the downregulation of long genes with CDK7 inhibition is likely a secondary effect of the generalized failure in transcription initiation efficacy following holoenzyme collapse, as longer transcripts require higher processivity. Regarding CDK9 inhibition, our previous work indicated that its inhibition affects long gene downregulation [31]. Here, our data displayed a similar trend, although the effect was less pronounced (**Suppl. Figure 3B**). Notably, under CDK12 inhibition, DNA repair pathways, particularly homologous recombination (HR), were markedly enriched among the downregulated genes (**Fig. 2D and Suppl. Figure 4**). It is noteworthy that most HR repair genes belong to the “long gene” category. This result indicates that the “condensation program” triggered by CDK12 loss progressively suppresses long gene transcription, ultimately depleting the DNA repair reservoir of cell.

### CDK12 deficiency induces progressive genome-wide chromatin compaction

Our results thus far indicate that CDK12 inhibition initiates a “chromosome condensation” program after 4 hours treatment. To determine whether the enrichment of condensation signatures observed at the transcriptional level translates into physical chromatin alterations, we evaluated chromatin accessibility under major transcriptional kinases inhibition using ATAC-seq. Given the potential time lag between the issuance of transcriptional instructions and the remodeling of large-scale 3D chromatin structures, we focused on a time-course ATAC-seq analysis (4h and 24h) to trace the physical trajectory of this process following CDK12 inhibition. Meta-gene analysis revealed a distinct, two-stage dynamic evolution (**Fig. 3A**). In the acute phase of CDK12 inhibition, chromatin accessibility signals near transcription start sites (TSS) dropped sharply to their lowest levels across all time points (**Fig. 3A**). This result reflects an “acute promoter closure” resulting from the loss of CDK12 activity. To exclude the possibility that this is a universal effect of transcriptional kinase inhibition, we compared these data with CDK7 (YKL-5-124) and CDK9 (NVP-2) inhibitions. While all treatments led to some degree of chromatin compaction, the effect was most pronounced under CDK12 inhibition (**Suppl. Figure 5**), highlighting the uniqueness of the TSS signal pattern induced by CDK12 inhibition. We hypothesized that the cellular response to CDK12 inhibition-mediated compaction might involve defensive compensatory mechanisms. However, the transcription factor (TF) motifs analysis revealed no compensatory activation of “rescue” factors such as E2F (**Fig. 3B**). Instead, we detected a broad loss of binding activity for general transcription factors, such as SP1 and KLF (**Fig. 3B**). This suggests that cells devoid of CDK12 activity bypass the “stress compensation” phase; rather, the acute shedding of general TFs, presumably together with histone modification changes, directly drives a large part of acute genome shutdown. As inhibition extended to 24 hours, although the mean TSS signal showed a slight recovery likely due to background noise from cellular stress, genome-wide peak analysis revealed a catastrophic loss of chromatin regions (**Fig. 3A**). These results can be validated by quantitative analysis. We identified over 33,000 significantly closed differentially accessible regions (DARs), a number far exceeding those observed at earlier time points (**Fig. 3C and Suppl. Figure 6)**. These results confirm that CDK12 inhibition leads to widespread chromatin compaction.

**Figure 3.**
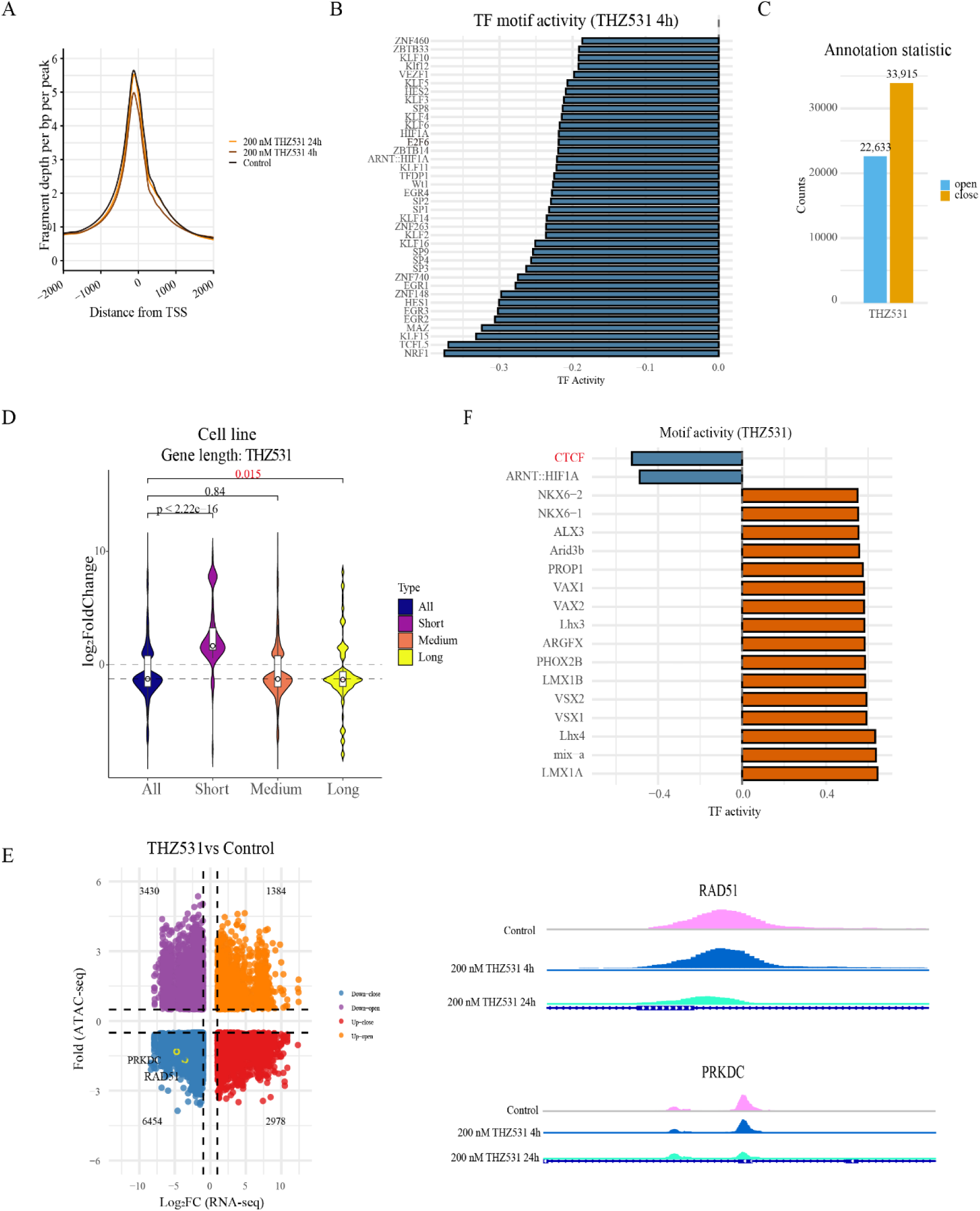
CDK12 deficiency triggers chromatin compaction and CTCF eviction. **A** Metagene plots of ATAC-seq signals centered on transcription start sites (TSS) in C4-2 cells treated with THZ531 for 4 h (acute phase) and 24 h (late phase), showing loss of chromatin accessibility. **B** Transcription factor (TF) motif analysis in regions losing accessibility at 4 h, showing loss of general TFs like SP1 and KLF. **C** Quantification of differentially accessible regions (DARs). Bar plot shows the number of significantly closed regions (FDR < 0.05) upon CDK12 inhibition (24h) compared to controls. **D** Integration of RNA-seq and ATAC-seq data. The violin plot shows that CDK12 inhibition suppresses the expression of long genes in DARs and promotes the expression of short genes in DARs. Significance was assessed using Student’s *t*-test. Gene length grading is consistent with Figure 2C. **E** Four-quadrant plot correlating changes in chromatin accessibility (ATAC-seq) and gene expression (RNA-seq) with CDK12 inhibition. Left: CDK12 inhibition primarily affects the downregulation of significantly close genes. A four-quadrant plot showing changes in chromatin accessibility (ATAC-seq) and gene expression (RNA-seq) after CDK12 inhibition (200 nM THZ531, 24h) reveals that significantly closed-down (6454) genes constitute the largest proportion. Right: ATAC-seq signal trajectory visualization shows that in closed genes under THZ531 treatment, the ATAC signal at the *PRKDC* and *RAD51* sites in long genes related to DNA repair remained unchanged after 4h treatment but were more closed after 24h treatment. **F** Motif analysis of TF binding sites in chromosome 24h after THZ531 (CDK12/13) treatment shows a significant enrichment of CTCF motif loss.

To confirm whether the physical densification caused by CDK12 inactivation directly drives long gene silencing, we integrated the ATAC-seq data with RNA-seq data. Gene length-dependent analysis of genes associated with significant DARs showed that long genes were significantly downregulated within the differential chromatin accessibility population (**Fig. 3D**). We further visualized the relationship between gene expression changes and chromatin accessibility using a four-quadrant plot (**Fig. 3E**). The results show that the majority of downregulated genes, including long DNA repair genes such as *PRKDC* and *RAD51*, fell into the third quadrant (Close-Down), characterized by both chromatin closure and transcriptional downregulation. *PRKDC* and *RAD51* participate in two core pathways of DNA double-strand break repair: non-homologous end joining (NHEJ) and homologous recombination (HR). Their relatively long gene length makes their transcription highly dependent on CDK12 to suppress premature intron polyadenylation (IPA) and maintain elongation persistence [14, 16]. Our IGV tracks further provided direct evidence of this spatiotemporal evolution. At the *PRKDC* and *RAD51* loci, chromatin open peaks showed only a minor reduction or maintenance after 4 hours of treatment compared to controls, confirming that the physical scaffold had not yet fully collapsed during the early instructional phase. However, by 24 hours, chromatin signals in these key regions had significantly diminished (**Fig. 3E**). These results demonstrate that the silencing of long gene transcription is explained, at least in part, by chromatin condensation. We hypothesized that the motif analysis could further explain the molecular driver(s) of this chromatin accessibility-change. At the 24-hour treatment, the binding activity of the chromatin architectural protein CTCF was significantly decreased (**Fig. 3F**). As CTCF serves as a critical anchor protein for maintaining topologically related domains (TADs) and chromatin loops [33, 34], its specific loss marks the complete failure of compensatory mechanisms and trigger the entry of chromatin into compaction phase. Our findings here, in particular the reduced CTCF binding activity detected when CDK12 is inhibited, are related to chromatin closure.

Subsequently, to investigate whether this outcome is a specific consequence of decrease in CDK12 activity, we employed CDK9 inhibition (NVP-2) as a mechanistic control. Surprisingly, gene length-dependent analysis of significant DARs under CDK9 inhibition revealed a state diametrically opposed to that of CDK12 inhibition (**Suppl. Figure 7A**). CDK9 inhibition induced a significant increase in chromatin accessibility (opening) across long gene regions (**Suppl. Figure 7A**). This phenotype is consistent with the RNA Pol II “pausing-stacking” mechanism induced by CDK9 inhibition [35], where accumulated protein complexes physically support open local chromatin. Furthermore, we compared TF motifs response profiles between the two treatments. Under CDK9 inhibition, we observed significant activation of the E2F family (e.g., E2F1), key regulators of the cell cycle [36] (**Suppl. Figure 7B**). We speculate that this E2F-mediated compensation, combined with the RNA Pol II pausing-stacking effect, successfully maintains or even expands chromatin openness at long gene loci, at least acutely after inhibition [37, 38]. This comparison provides compelling evidence that chromatin compaction at long gene loci is not a generic consequence of elongation blockade, but rather a result of inhibition induced by a specific transcriptional kinase, CDK12/13.

In summary, these data delineate a unique chromatin landscape induced by decrease in CDK12 activity, evolving from a general lack of TFs in the early phase to a decrease in CTCF binding activity in the late phase resulting in genome-wide architectural compaction. This chromatin blockade constitutes the structural basis for the transcriptional silencing of long genes, particularly DNA repair genes. Standing in stark contrast to the RNA Pol II pausing effect caused by CDK9 inhibition, these findings establish CDK12 not merely as a transcriptional elongation kinase, but as a critical “gatekeeper” maintaining the topological integrity of long genes.

### *In vitro* chromatin compaction correlates with long gene silencing in *CDK12*-mutant prostate tumors

To further assess the clinical relevance of *CDK12* deficiency-induced chromatin compaction in patient tumors, we performed a multi-omics integrative analysis. We combined our *in vitro* ATAC-seq data (CDK12 inhibition) with expression data from the TCGA database derived from metastatic and primary PC patients harboring *CDK12* truncating mutations (**Fig. 4A and Suppl. Figure 8**). Quantitative analysis of the four-quadrant distribution revealed that genes associated with a loss of chromatin accessibility dominated the landscape in both primary and metastatic samples, accounting for 67.2% and 73.3% of the total (red plus blue regions), respectively (**Fig. 4B left**). Strikingly, in metastatic samples, the “Down-close” sub-group, characterized by concomitant chromatin closure and transcriptional downregulation, was the most significantly enriched, representing 42.1% of the analyzed genes (**Fig. 4B**). Further statistical analysis clarified the link between chromatin compaction and gene downregulated. We found that among all downregulated genes (blue and purple quadrants), over 80% in metastatic samples were accompanied by chromatin closure, compared to 56.9% in primary samples (**Fig. 4B, right**). These results suggest that chromatin compaction resulting from *CDK12* deficiency is the dominant force driving gene downregulated in clinical PC samples, and this characteristic manifests more severely in the metastatic subtype.

**Figure 4.**
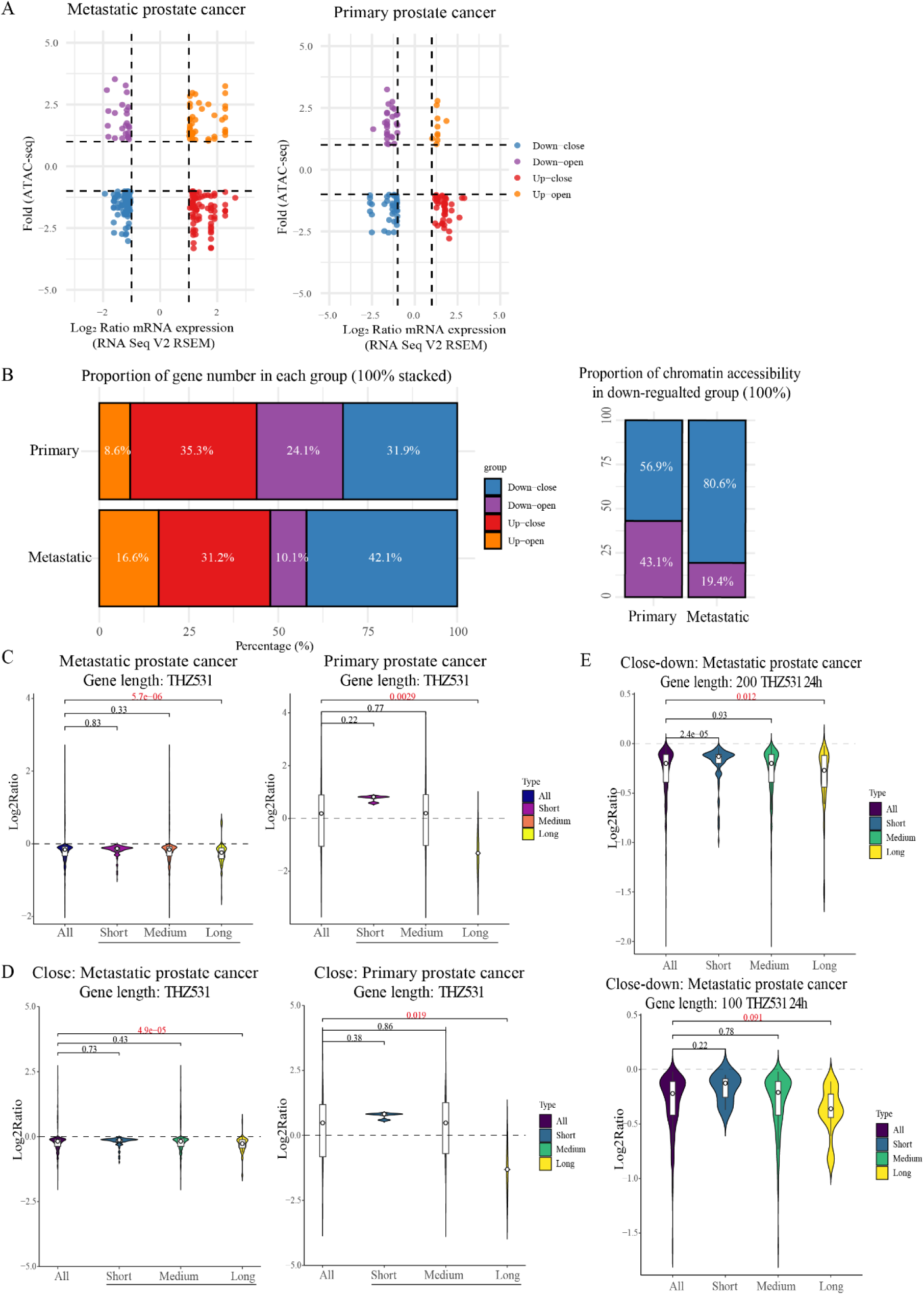
Chromatin compaction is associated with long gene silencing in prostate patient tumors with *CDK12* truncating mutations. **A** The four-quadrant diagram illustrates the distribution of DARs data integrating *in vitro* CDK12 inactivation and differential expression profiles of CDK12 truncated mutations in TCGA prostate cancer patients (metastatic and primary). Patient data is derived from the TCGA database. **B** Statistical analysis of DEGs and DARs in prostate *CDK12*-mutant patient samples. Left: Percentage of DEGs based on DARs (in vitro) and the number of *CDK12*-mutant primary and metastatic PC patients. Right: Proportion of genes with chromatin changes among downregulated genes in metastatic and primary prostate tumors. **C** Gene length-dependent expression analysis of TCGA prostate patient tumors harboring CDK12 truncating mutations. The violin plots show that *CDK12* truncated mutations significantly downregulated the expression of long genes in both metastatic and primary prostate patient tumors. **D** In metastatic and primary prostate cancer samples, long genes located in significant chromatin-closed regions were significantly downregulated in *CDK12* truncated mutation patient samples. **E** Analysis of a significant subset of *CDK12*-inactivated “close-down” genes in metastatic samples highlighted specific repression of long genes.

To verify whether this *CDK12* inactivation-driven chromatin densification directly precipitates long gene downregulation in clinical context, we analyzed metastatic and primary PC patients with CDK12 truncated mutations separately. First, direct gene length-dependent analysis of the *CDK12*-mutant patient transcriptomes mirrored our *in vitro* findings, confirming that the specific downregulation of long genes is a feature of this tumor subtype (**Fig. 4C**). Subsequently, by mapping the patient-derived downregulated gene sets back to our epigenomics data, we found that long genes located within chromatin closed regions displayed significant length-dependent downregulation (**Fig. 4D**). Finally, focusing specifically on the core “Close-down” gene set, metastatic samples again exhibited distinct transcriptional defects in long genes (**Fig. 4E**).

Collectively, these findings show that in clinical prostate patients tumor samples, particularly within the metastatic subtype, the chromatin closure induced by *CDK12* inactivation functions as a “structural blockade” specifically blocking and downregulating long genes.

### Proteomic analysis reveals specific recruitment of adaptive remodeling factors upon CDK12 inhibition

Despite the dual pressures of chromatin compaction and transcriptional blockade, cells did not undergo immediate death. This suggests that in the context of CDK12 deficiency, the transcriptional machinery may initiate an adaptation to maintain basal survival and even confer growth-advantage. To identify the key factors maintaining this “adaptive balance” and to understand how cells respond to the chromatin stress imposed by decrease in CDK12 activity, we utilized our 5-EU metabolic labeling strategy (**Fig. 1**). This approach captures the “active” protein interactome that associates with the transcription machinery during the imposed stress. We treated the cells with CDK12/13 inhibitor THZ531 for 4 hours, followed by a 5-EU pulse for the last 10 minutes of treatment, and then collected samples for MS. To assess the potential common factors for transcriptional stress and specifically CDK12-dependent, we performed parallel evaluations using inhibitors of other major transcriptional kinases, CDK7 (YKL-5-124) and CDK9 (NVP-2). Statistical analysis of differential protein enrichment indicated that among all treatment groups, CDK12 inhibition triggered the most profound proteomic remodeling, followed by CDK7 and CDK9 inhibition (**Fig. 5A**). Our data also showed that while kinase inhibition led to the decrease from actively transcribing RNA Pol II of most transcription-dependent proteins, reflecting defects to the transcriptional machinery, a subset of proteins in each group exhibited significantly increased association (**Fig. 5A**). These proteins, increased against the general trend of suppression, imply that following kinase inhibition, specific factors are recruited to remodel RNA Pol II complexes, and constitute part of an “adaptive response” to transcriptional perturbation.

**Figure 5.**
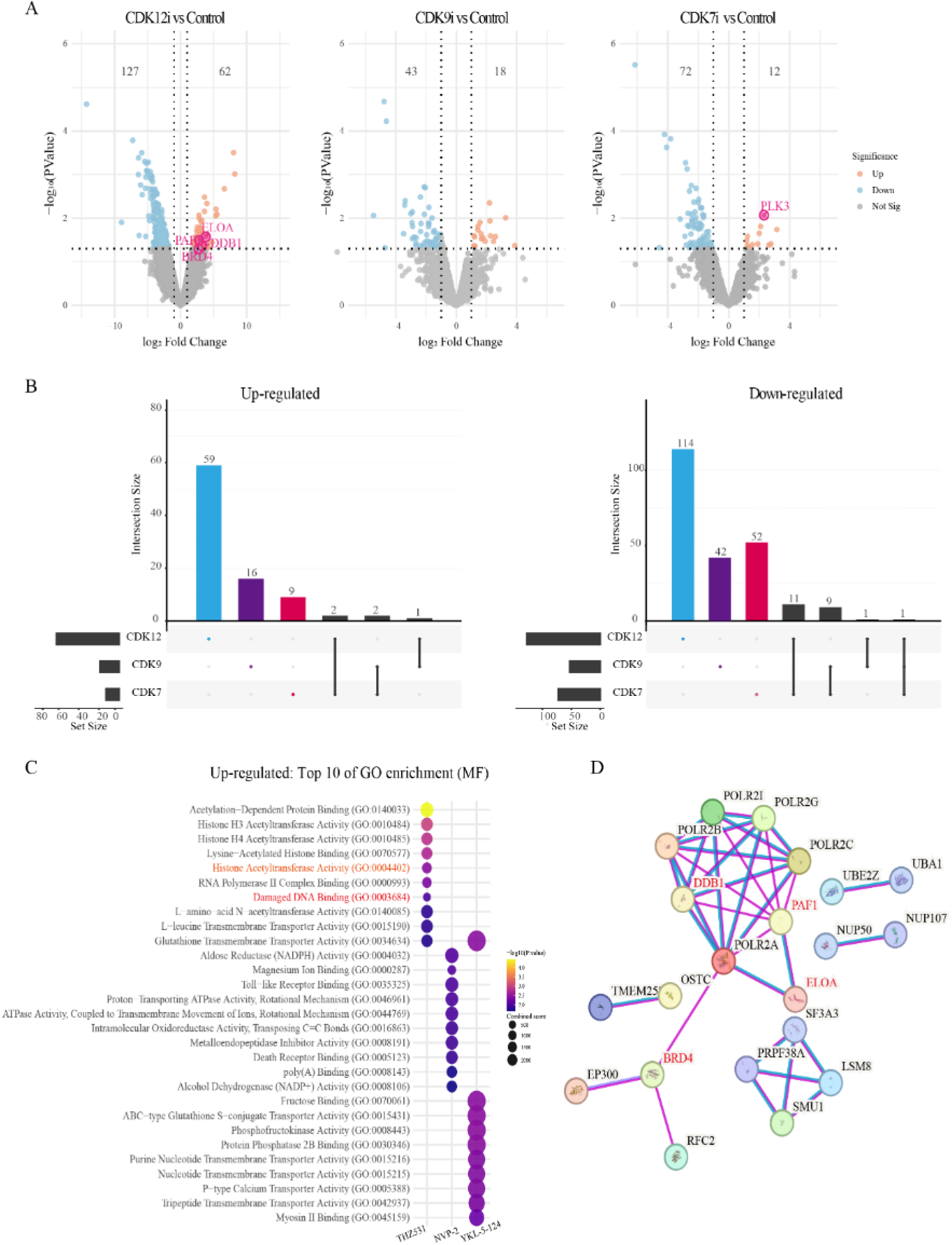
5-EU-labeled proteomics mapping reveals adaptive recruitment to RNA Pol II after CDK12/13 (THZ531) inhibition. **A** Volcano plot of RNA Pol II-interacting proteins in C4-2 cells treated with 150 nM THZ531 (CDK12/13i), 20 nM NVP-2 (CDK9i) and 500 nM YKL-5-124 (CDK7i) for 4 h. The results show that 127, 43 and 72 proteins are significantly enriched under CDK12i, CDK9i and CDK7i, respectively, and 62, 18 and 12 proteins are significantly decreased (p < 0.05 and log2FC ± 1). **B** Upset plots compare the distribution of proteins that are significantly lost and gained after inhibition of CDK7, CDK9, or CDK12 under the 5-EU metabolic labeling method. **C** The dotplot shows the GO molecular function enrichment analysis of proteins that were significantly increased after inhibition of CDK7, CDK9, and CDK12. CDK12 inhibition specifically upregulated and enriched the “Damaged DNA Binding” and “Histone Acetylation” pathways, while CDK7i and CDK9i do not. **D** The protein-protein interaction (PPI) network (STRING database) of the adaptive complex formed around RNA Pol II, which exhibits significant increased interaction with RNA Pol II upon CDK12 inhibition, is centered on DDB1, BRD4, PAF1, and ELOA.

Subsequent upset plot analysis not only visualized the distribution of differentially expressed proteins but also revealed distinct recruitment signatures for each kinase inhibition (**Fig. 5B**). CDK7 inhibition specifically enriched the cell cycle regulator *PLK3* (**Fig. 5A**). In contrast to the PLK3 enrichment induced by CDK7 inhibition, CDK12 inhibition induced the enrichment of a specific set of chromatin remodelers and transcriptional co-factors. Notably, Pol II associated factor 1 (PAF1), Damage-specific DNA binding protein 1 (DDB1), Bromodomain containing 4 (BRD4), and ELOA were significantly enriched at the protein level (**Fig. 5A and B**). DDB1, as a core subunit of the CUL4-DDB1 ubiquitin ligase, may participate in the clearance of stalled RNA Pol II or the remodeling of the compacted chromatin [39]. BRD4 and PAF1 are likely recruited to assist transcription elongation or to maintain enhancer function, attempting to overcome the elongation barrier imposed by CDK12 deficiency. This hypothesis is supported by previous studies demonstrating that BRD4 recruits the elongation factor positive transcription elongation factor b (P-TEFb) to the RNA Pol II to stimulate elongation [40, 41]. Meanwhile, the PAF1 complex has been reported to directly promote RNA Pol II pause release by serving as a scaffold for elongation factors and stimulating the catalytic activity of the CDK12 and cyclin partner, ensuring efficient transcription of long genes [42]. Notably, CDK9 inhibition did not induce enrichment of these factors, further confirming that this adaptive response is specific to the unique remodeling triggered by CDK12 inhibition.

To systematically investigate whether these proteins with enhanced interaction with RNA polymerase II constitute a coordinated functional program, we performed functional enrichment analysis. GO molecular function (MF) analysis revealed that CDK12 inhibition specifically led to the enrichment of proteins related to “Damaged DNA Binding” and “Histone Acetylation” pathways that were absent in the CDK7 or CDK9 inhibition treatments (**Fig. 5C**). Furthermore, reactome pathway analysis indicated that CDK12 inhibition activated “DNA Damage Bypass” pathways (**Suppl. Figure 9**). This implies that in the absence of CDK12, which halts transcription of long genes including repair genes, cells may resort to these error-prone mechanisms like DDB1-mediated pathways to bypass barriers on chromatin and sustain survival. These results align closely with our earlier findings of chromatin compaction, indicating that cells attempt to alleviate structural blockaded genomes by recruiting DNA damage recognition factors (e.g., DDB1) and epigenetic regulators (e.g., BRD4).

We hypothesized that CDK12 inhibition leads to an adaptive remodeling of the RNA Pol II complex. To verify this, we imported the proteins significantly enriched during CDK12 inhibition into the STRING database to construct a Protein-Protein Interaction (PPI) network centered on RNA Pol II. The results showed that DDB1 (repair/remodeling), BRD4 (epigenetic reading), PAF1 (elongation assistance), and ELOA clustered around the Pol II core, displaying high-confidence interactions (**Fig. 5D**). This result strongly demonstrates that these factors are recruited in close proximity to the stalled RNA Pol II complex, forming a putative adaptive complex. In summary, this remodeling complex may represent a survival response employed by tumor cells facing the chromatin condensation caused by decrease in CDK12 activity, thereby exposing potential therapeutic targets.

### Targeting the adaptive factor PAF1 induces selective combinatorial lethality with CDK12/13 inhibition

Based on the RNA Pol II adaptive interactome landscape we identified under CDK12 inhibition, we hypothesized that these recruited proteins constitute a critical buffering mechanism employed by cancer cells to mitigate the stress of CDK12 deficiency. Consequently, severing this “stress-survival” pathway could block cellular escape routes and induce combinatorial lethality.

Before evaluating this combinatorial strategy, we first assessed the sensitivity of multiple prostate cancer cell lines (LNCaP, C4-2, 22RV1) and cell line derived from normal prostate epithelia (PNT1) to transcriptional CDK inhibitors (CDK7, CDK9, CDK12). Cell viability assays utilizing the same inhibitor concentrations used for proteomic enrichment revealed that cancer cells exhibited preferential susceptibility to the disruption of transcriptional CDK activity, whereas normal PNT1 cells displayed relative resistance to inhibitors targeting these kinases (**Fig. 6A**). The resulting heatmap provides a global view of the selectivity of these inhibitors in prostate cancer cells. This distinct “therapeutic window” suggests that combining low-dose CDK12 inhibitors with agents targeting adaptive factors could specifically kill cancer cells while sparing normal cells.

**Figure 6.**
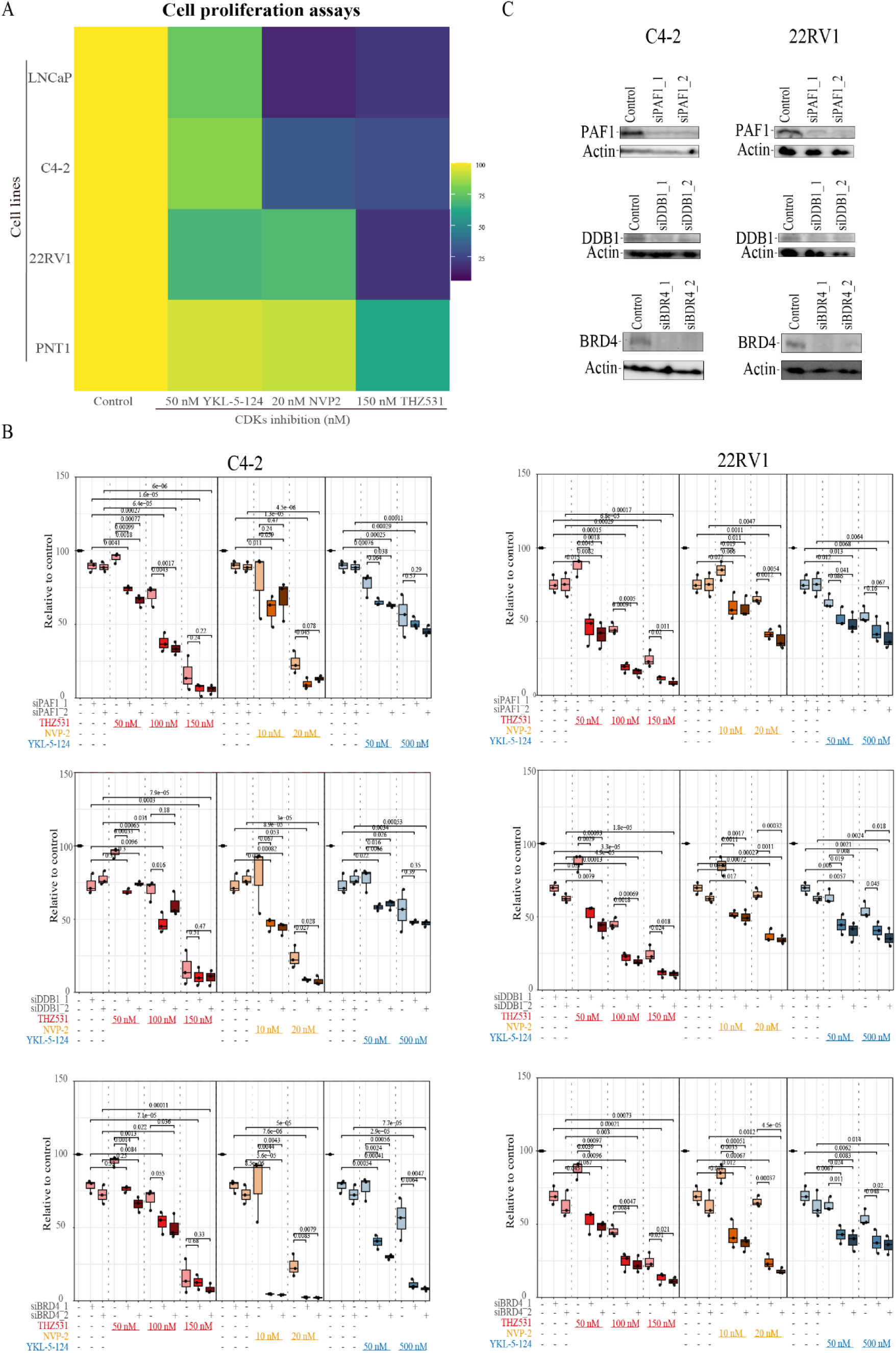
Targeting the adaptive factor PAF1 induces combinatorial anti-proliferative effects in prostate cancer cells in the context of CDK12 inhibition. **A** Prostate cancer cells depend on major transcription-associated cyclin-dependent kinases (CDK7, CDK9 and CDK12), whereas normal prostate cells show relative resistance to inhibitors targeting these kinases. Each cell line was cultured for 24 hours before treatment and then treated for 4 days in 50 nM YKL-5-124, 20 nM NVP-2 and 150 nM THZ531, respectively, and their cell viability was assessed using CellTiterGlo. The data is from three biological replicates, each having three technical replicates. **B** The results show that in the C4-2 cell line, PAF1 knockdown produced significant combined decrease in viability only with CDK12 inhibition. DDB1 knockdown produced significant combined decrease in viability with CDK12 and CDK9 inhibition, but not with CDK7 inhibition. BRD4 knockdown produced significant combined decrease in viability with CDK12, CDK9, and CDK7 inhibition. In the 22RV1 cell line, DDB1 and BRD4 knockdown produced combined decrease in viability with CDK12, CDK9, and CDK7 inhibition, but PAF1 knockdown produced significant combined lethality only with CDK12 inhibition. Cell viability after 4 days of DDB1, BRD4, and PAF1 knockdown with CDK7, CDK9, and CDK12 inhibitor treatments for the last three days. Data were obtained from three biological replicates (each containing three technical replicates), and the standard error of the mean was calculated. Student’s *t*-test was used to assess significance. **C** The knockdown of DDB1, BRD4, and PAF1 was confirmed by western blotting in castration-resistant prostate cancer cell lines C4-2 and 22RV1 (n=2).

To distinguish true functional vulnerabilities from the list of physically enriched candidates (DDB1, BRD4, PAF1, ELOA, PLK3), we performed an siRNA-based screen to evaluate cell viability in combination with different inhibitors. Notably, although ELOA and PLK3 displayed increased interaction with RNA Pol II under stress (THZ531 and YKL-4-125) (**Fig. 5**), their knockdown failed to induce significant combinatorial lethality in the tested cell lines (**Supplementary Fig. 10**). This indicates that not all factors recruited to RNA Pol II assume a core survival function, highlighting the necessity of functional validation.

In stark contrast, DDB1 knockdown (siDDB1) significant combinatorial anti-proliferative effects with CDK12/13 inhibitor THZ531 in both 22RV1 and C4-2 cells (**Fig. 6B**). To confirm the correlation between phenotype and protein levels, we validated the effective knockdown of DDB1, BRD4, and PAF1 in these core models via Western blotting (**Fig. 6C**). Although knockdown of other candidate adaptive factors, such as BRD4, also demonstrated significant sensitization, validating the general feasibility of the “targeting chromatin adaptive mechanisms”-strategy, PAF1 and DDB1, especially PAF1, showed unique cell type- and kinase-specificity. While BRD4 is an effective sensitization target, it acts as a general super-enhancer regulator that stimulates global transcription via CDK9 recruitment [43, 44]. Its function is not specifically confined to alleviating the stress caused by CDK12/13 inhibition. Conversely, PAF1 and DDB1 knockdown induced a more complete and potent combinatorial lethal effect than BRD4 knockdown in combination with CDK12/13 inhibitor (e.g., C4-2) (**Fig. 6B**).

Crucially, this dependency on PAF1 and DDB1 exhibited high kinase- and cancer cell line-specificity. First, regarding kinase specificity, DDB1 knockdown did not elicit equivalent combinatorial lethality when combined with CDK7 (YKL-5-124) inhibitor in C4-2 cell line (**Fig. 6B**). This demonstrates that DDB1 addiction is not a generic response to transcriptional interference. Notably, DDB1 depletion conferred sensitivity not only to CDK12 inhibition but also to CDK9 inhibition. This suggests that DDB1 may safeguard against broad transcriptional elongation stress or genomic instability mechanisms shared by the CDK9 and CDK12/13 pathways. In sharp contrast, PAF1 exhibited strict specificity for CDK12/13. PAF1 knockdown sensitized all prostate cancer cell lines to CDK12/13 inhibition, it did not induce significant combinatorial lethality under conditions of CDK7 (in C4-2 and 22RV1) or CDK9 (in C4-2) inhibition (**Figure 6B and Suppl. Figure 10A**). This striking contrast demonstrates that, unlike DDB1, PAF1 addiction is a vulnerability uniquely conferred by the specific stress triggered by CDK12 loss.

Next, to validate the broad applicability of this strategy, we evaluated other prostate cancer cells (LNCaP) and cell line derived from normal prostate epithelia (PNT1). Consistent with the results in 22RV1 and C4-2, LNCaP cells showed significant sensitivity to the combination of THZ531 with either siPAF1 or siDDB1 (**Suppl. Figure 11A**). Importantly, in normal PNT1 cells, the same combinatorial treatment did not result in significant decrease in cell viability (**Supplementary Fig. 11B**). These data suggest that normal cells can tolerate the transient loss of PAF1 or DDB1 when CDK12 is inhibited.

Collectively, our data identify a potential therapeutic window: while DDB1 is a broad-spectrum extended stress target (CDK9 and CDK12) and does not have effects on normal cells, PAF1 is a promising therapeutic target for *CDK12*-deficient tumors. Targeting PAF1 exploits the cancer-specific adaptive stress mechanism to induce potential synthetic lethality.

## Discussion

In this study, by integrating the mapping of dynamic RNA Pol II interactome strategy with multi-omics profiling, we show a non-classical function of CDK12 as a regulator of chromatin topology. We show that CDK12 inhibition induces a unique “structural blockade” characterized by chromatin compaction (**Figs. 2B and 3A**), which physically precludes the transcription of long genes, particularly those governing DNA repair (**Fig. 2C and 3D**). Furthermore, we identify a specific adaptive survival mechanism where PC cells recruit a DDB1- and PAF1-nucleated complex to RNA Pol II to navigate this stress (**Figs. 5A and 5D**). This finding further confirms that targeting PAF1 specifically induces combinatorial lethality in CDK12-inhibition prostate cancer cells and constitutes a potential therapeutic target (**Fig. 6B**).

Our data challenge the traditional view that transcriptional kinases merely regulate the enzymatic rate of RNA Pol II. We found that inhibition of the major transcriptional regulatory kinases CDK7, CDK9, or CDK12 all led to chromatin accessibility compression (**Fig. 3A**), reflecting a general coupling between transcriptional activity and open chromatin [19, 45]. Interestingly, however, only the elongation kinase CDK12 inhibition initially provided the signal signature for chromosome compression (**Fig. 2B**). This implies that the nature of the chromosome accessibility changes resulting from the inhibition of these transcriptional kinases is fundamentally different. As a general transcription initiation kinase, CDK7 inhibition primarily leads to the downregulation of holoenzyme complex-related genes (**Supplementary Fig. 2B**) [30], resulting in a “passive collapse” following the breakdown of the transcriptional machinery. Similarly, while CDK9 inhibition also restricted chromatin accessibility, this likely derives from distinct molecular events. As the core catalytic subunit of the P-TEFb, CDK9 functions primarily by phosphorylating the Ser2 residue of the RNA Pol II CTD as well as the negative elongation factors NELF and DSIF, thereby releasing promoter-proximal pausing and driving productive elongation [46, 47]. Consequently, CDK9 inhibition directly blocks this key release step, “freezing” RNA Pol II in a promoter-proximal paused state [35]. These factors may lead to a chromatin landscape consistent with functional elongation arrest of the transcription machine.

In a marked contrast, CDK12 inhibition drives the genome into a state of structural collapse, characterized by the profound chromatin compaction observed in our study (**Fig. 3A and Supplementary Fig. 5**). This phenomenon is intrinsically linked to the dual function of CDK12 as an important regulator of both transcription elongation and genomic stability [12, 16, 48].

In prostate cancer, *CDK12* truncating mutations define an aggressive subtype, characterized by genomic instability that is attributed to the loss of expression of DNA repair genes [14]. However, our findings reveal a more fundamental possible mechanism, in which *CDK12* loss extends beyond the mere limitation of kinase activity to trigger decrease in the binding force between the architectural protein CTCF and chromatin (**Fig. 3F**). This implies a topological collapse, wherein the abrogation of *CDK12*-dependent phosphorylation compromises the maintenance of chromatin loops or topologically associating domains (TADs) [34, 49]. The erosion of these boundaries facilitates the aberrant spreading of heterochromatin [50], establishing a physical barrier that is particularly harmful to the transcriptional processivity required for long genes (**Figs. 3D and 3E**) [16]. In practice, it is likely that both decreased expression of DNA repair genes and chromatin compaction contribute to the phenotype of cells impacted by CDK12 inactivation.

This “chromatin roadblock” brought about through chromatin compaction, bears potential clinical indication. We observed an intrinsic coupling between the regions of closed chromatin (ATAC-seq) and downregulation of long genes in patients with metastatic prostate cancer (**Figs. 4B and 4D**). These physical constraints offer a structural rationale for the TDP characteristic of *CDK12*-mutated tumors [14]. In practice, we propose that the accumulation of unresolved topological stress within long genes renders these loci susceptible to catastrophic genomic instability [51, 52]. Unlike single-gene mutations, this concurrently silences multiple DNA repair pathways through physical compaction (**Figs. 2D and 3D**), and this length-dependent downregulation was confirmed in a cohort of prostate patients with *CDK12* truncated mutations (**Fig. 4C**). Notably, this signature of “compaction-driven silencing” is markedly more pronounced in metastatic samples compared to primary tumors (**Figs. 4C-E**). This likely reflects the intrinsic biology of mCRPC, where advanced tumors typically exhibit elevated basal replication stress [53–55]. Crucially, while this transcriptional defect imposes stress, the specific silencing of long DNA repair genes fuels genomic instability, specifically the TDP, which acts as a driver of tumor heterogeneity and aggressiveness [14, 15]. However, this mutant state renders tumor cells acutely sensitized to the additional topological constraints imposed by *CDK12* loss, such as R-loop accumulation and transcription-replication conflicts, thereby exacerbating the transcriptional collapse of long genes [55].

Crucially, our proteomic analysis reveals how cells adapt to decrease in CDK12 activity. We observed that PAF1, DDB1 and BRD4 are specifically recruited to RNA Pol II following CDK12/13 inhibition (**Figs. 5A and 5B**) and cell cycle regulator PLK3 was specifically enriched in CDK7 inhibition (**Fig. 5A**). The recruitment of PLK3 is consistent with the roles of both CDK7 and PLK3 in regulating progression through the cell cycle [56, 57]. Our previous studies have shown that CDK12 inhibition leads to rapid decrease in histone H3 acetylation, which is likely a direct consequence of CDK12 inhibition causing the ATAC acetyltransferase complex to detach from RNA Pol II [31]. Given that histone hypoacetylation is a hallmark precursor to chromatin compaction and transcriptional silencing, this suggests that the recruitment of DDB1 is not coincidental. Rather, the cell perceives the resulting repressive chromatin environment not only as transcriptional suppression, but also as a form of urgent architectural stress or stalling necessitating resolution.

While the recruitment of the epigenetic reader BRD4 reflects a generic attempt to re-initiate transcriptional elongation via P-TEFb (CDK9) recruitment [58], our combinational inhibition screens identified DDB1 as a critical vulnerability required for survival under CDK12/13 inhibition (**Fig. 6B**). Unlike BRD4, which functions as a global transcriptional regulator, DDB1 serves as a core component of the CRL4 ubiquitin ligase complex, specialized in sensing DNA damage and stalled replication forks [59, 60]. Consequently, its specific recruitment likely represents an effort by the cell to clear stalled polymerases entrapped by compacted chromatin or to remodel local topology. This adaptive attempt possibly aimed at resolving transcription-replication conflicts (TRCs) or clearing stationary polymerases trapped in the dense chromatin [61, 62]. Although we have established DDB1 as a key executor in this process, the precise molecular mechanisms warrant further elucidation.

While the recruitment of DDB1 reflects an attempt to clear stalled polymerases from the compacted chromatin, our combinational inhibition screens identified PAF1 as a more specific vulnerability required for survival under CDK12 inhibition (**Fig. 6B**). The PAF1 complex (PAF1C) is known to play a critical role in promoting RNA Pol II pause-release [42]. Unlike DDB1, which represents a broad-spectrum response factor to elongation stress (sensitive to both CDK9 and CDK12 inhibition), PAF1 dependency exhibits selective combination lethality effect for CDK12 inhibition. Thus, PAF1 knockdown effectively converts this adaptive response into a lethal trap. The “therapeutic window” we observed, in which cells could tolerate PAF1 depletion in the absence of the basal chromatin pressure associated with *CDK12* deficiency (**Figs. 6A, 6B and Suppl. Fig. 11B**), further highlights the translational potential of this strategy. Furthermore, given the association of this dependency with high replication stress, future investigations should determine whether this synthetic lethal strategy extends to other tumor types harboring analogous genomic features.

## Conclusion

We extend the role of CDK12 from a regulator of transcription elongation to a gatekeeper of chromatin topology. Our comprehensive mapping of the actively transcribing RNA Pol II interactome, chromatin accessibility and transcriptome reveal that CDK12 inhibition precipitates a unique structural blockade, characterized by the loss of CTCF and chromatin compaction. This physical constraint serves as the mechanistic driver for the silencing of long genes, distinguishing *CDK12* deficiency from the transcriptional pausing induced by CDK9 inhibition. Furthermore, our work shows a specific adaptive response enrichment wherein PC cells actively recruit a DDB1- and PAF1-nucleated complex to RNA Pol II to navigate this architectural stress. This recruitment likely serves as a quality control step to clear stalled polymerases entrapped by the compacted chromatin. Consequently, we propose that this dependency creates a therapeutic window, by targeting PAF1 to exploit cancer-specific adaptation to induce synthetic lethality in *CDK12* mutant cells. Our findings provide a molecular rationale for combining PAF1 inhibition with *CDK12* inactivation, offering a promising actionable avenue to selectively eliminate aggressive, genomic instability-driven prostate tumors.

## Materials and methods

### Cell culture, proliferation assays and western blotting

C4-2, 22RV1 and LNCaP cell lines were obtained from the American Tissue Culture Collection, and the PNT1 cell line was acquired from Sigma. These cell lines were maintained in RPMI medium supplemented with 10 % fetal bovine serum (FBS). Transfection of Ambion® Silencer Select siRNAs against DDB1 (Assay IDs s3980 and s3981), BRD4 (Assay IDs s23901 and s23903), PAF1 (Assay IDs s29267 and s29268), PLK3 (Assay IDs s3245 and s3246), and ELOA (Assay IDs s224710 and s13859) were achieved using RNAiMax (Invitrogen™ Lipofectamine™ RNAiMAX Transfection Reagent, Catalog number: 13778150). For viability assays, knockdown was performed for 24 hours, after which the indicated treatments were added for 4 days. Next, CellTiter-Glo® 2.0 assay (Promega) was used according to the manufacturer’s instructions in technical triplicates in 384-well plates and the data presented is from three biological replicates.

For western blot experiments, cells were treated as indicated in the figure-legends. Cell lysates for western blot were prepared as previously described (except no sonication) [63]. Protease inhibitor cocktail (catalog number, HY-K0010) and phosphatase inhibitor (catalog number, HY-K0021) were purchased from MCE. Antibodies used are from ThermoFisher Scientific: DDB1 (2B12D1) and PAF1 (A300-172A); Santa Cruz Biotechnology: RNA Pol II (sc-56567), GAPDH (sc-47724); Proteintech: BRD4 (28486-1-AP) and from Abcam: Actin (ab49900). Compounds were obtained from MedChemExpress.

### RNA polymerase II immunoprecipitation and mass spectrometry

For RNA Pol II immunoprecipitation and mass spectrometry (MS) experiments, we combined two antibodies (both from Santa Cruz Biotechnology): CTD4H8 (sc-47701) and 8WG16 (sc-56767). Before collection, C4-2 cells were grown for two days without any treatment. Cells were washed with cold PBS, collected by centrifugation and solubilized into lysis buffer (50 mM Tris-HCl, pH 7.4, 2 mM EDTA, and 300 mM NaCl) with protease and phosphatase inhibitors and incubated for 30 minutes on rotation in the cold room. After centrifugation, the NaCl-concentration of the samples was adjusted to 100 mM. Samples were pre-cleared with 2 micrograms of IgG (Santa Cruz Biotechnology: sc-2025) and protein A/G Magnetic Beads (MedChemExpress: HY-K0202), after which overnight immunoprecipitation was performed using either Pol II-specific antibodies or the IgG control. The next day, protein A/G Magnetic Beads were added to samples (2 hours incubation) and samples were washed three times with washing buffer (50 mM Tris-HCl, pH 7.4, 2 mM EDTA, and 100 mM NaCl) and once with PBS. Mass spectrometry-analysis was purchased as service from University of Turku.

### Metabolic labeling of RNA using 5-ethynyl uridine (5-EU) and mass spectrometry

C4-2 cells were grown for two days, after which they were treated for 4 hours with 500 nM YKL-5-124, 20 nM NVP2, 150 nM THZ531, and without treatment. During the last 10 minutes of treatment, cells were labeled with 1 mM 5-EU. The groups not treated with inhibitors included both 5-EU labeled and unlabeled cells. Cells were fixed by 1% formaldehyde for 15 minutes and the reaction was quenched by treatment with 125 mM Glycine for 10 minutes after labelling. Cells were washed with cold PBS and collected by centrifugation at 4°C for 5 minutes. After centrifugation, cells were resuspended with permeabilization buffer (0.5% Triton X-100) and incubated at room temperature for 30 minutes. Cells were collected and incubated with “click-reaction” buffer at room temperature for 30 minutes after permeabilization. The click-reaction buffer was prepared according to the instructions of the Invitrogen manufacturer’s Click-iT® Cell Reaction Buffer Kit (catalog number, C10269) and Biotin Azide (PEG4 carboxamide-6-Azidohexanyl Biotin, catalog number B10184). After incubation, cells were collected by centrifugation (2000 rpm) and washed with cold 0.5% BSA. Cells were then lysed with lysis buffer (1% SDS in 50 mM Tris pH 8.0 with 1x protease inhibitor) for at least 30 minutes and sonicated (MSE Soniprep 150, settings three amplitude microns for 5 times, 20 seconds on, 40 seconds off on ice). The lysate was then collected by centrifugation with full speed for 10 minutes and diluted 1:10 using cold PBS. Streptavidin MagneSphere® Paramagnetic Particles (Z5481) were purchased from the Promega and had been washed three times with lysis buffer and once with PBS, were added to the lysate and incubated overnight at 4°C. The next day, streptavidin beads were washed three times with lysis buffer and once with PBS. Mass spectrometry-analysis was purchased as service from University of Turku. Western blotting was used to analyze untreated 5-EU labelled and unlabeled samples.

### SLAM-seq and ATAC-seq

C4-2 cells were grown in 6-well plates for one day. The next day, cells were untreated and treated with 500 nM YKL-5-124, 20 nM NVP2, 200 nM THZ531 for 4 hours, and 100 nM and 200 nM THZ531 for 24 hours. In the last 10 minutes before 4 hours and 24 hours, cells were labelled with 1 mM 4-thiouridine (4sU, obtained from ThermoFisher Scientific). RNA was purified using the Amersham RNAspin Mini Kit (catalog number 25050071) according to manufacturer’s instructions except 0.1 mM DTT was included in all the buffers. Purified RNA was alkylated as previously described [18, 64]. Library preparation and sequencing were bought as a service from Single Cell Genomics Core, University of Eastern Finland. In brief, libraries were prepared with the QuantSeq 3′ mRNA-Seq V2 kit with UDI and sequencing was performed using SR100 High Output sequencing on Illumina NextSeq 2000.

For ATAC-seq, C42 cells was grown in 6-well plate for 1 day. The next day, cells were treated with the same conditions as in the SLAM-seq experiment. Sample collection and library preparation of ATAC-seq were performed using the Hyperactive ATAC-Seq Library Prep Kit for Illumina (TD711) and TruePrep Index Kit V2 for Illumina (TD202) according to the manufacturer’s instructions. Subsequently, the sequencing was purchased as service from the BGI using PE100bp on the DNBseq platform.

### Bioinformatics data analysis

All analysis and visualizations are performed using the following main methods and tools, employing Shell, R, and Python. The SLAM-seq data analysis is as shown in our previous analysis method [18]. Gene length analysis also followed the gene length sources and classification criteria we used previously [18, 65], which can be summarized as short: <1 kb, medium: 1–15 kb, and long: >15 kb. ATAC-seq data were filtered using the same standards and methods as SLAM-seq data and aligned to the hg38 reference genome using bowtie2 [66]. Deeptools [67], HOMER [68], and DESeq2 [69] were used to assess chromosome accessibility signals, perform differential accessibility analysis and annotation, and visualize by removing mitochondria and duplications. The JASPAR [70, 71] database was used for transcription factor (TF) motif analysis. Enrichment analysis was performed using Enrichr [72] for GO [73] and KEGG [74].

Differentially expressed genes (DEGs) from RNA-seq were analyzed in conjunction with differentially accessible regions (DARs). Similarly, we used the cBioPortal to identify the prostate cancer patients with *CDK12* truncating mutations from the TCGA-PRAD dataset (metastatic and primary prostate patients) [75, 76]. Subsequently, gene length analysis was performed on the expression data, and this data was integrated with DARs data, followed by another gene length analysis.

For mass spectrometry data, differential abundance protein analysis was performed by fitting linear models, using weighted least squares to estimate differential protein abundance, and p-values were adjusted for multiple testing using the Benjamini–Hochberg procedure to identify significantly changed proteins (p< 0.05 and abs (Log(2FC))>1). STRING [77] database was used to perform PPI network analysis.

## Availability of data and materials

All the SLAM-seq data and ATAC-seq have been deposited to NCBI with the following accession numbers: PRJNAXXXXXXX and PRJNAXXXXXXX. Any additional information in this paper is available from the contacts upon reasonable request.

## Supplementary Information

This article contains one supplementary file, which contains supplementary figures 1-11 and one supplementary table 1.

## Acknowledgements

The authors are grateful to CSC—IT Center for Science, Finland, for the computational resources. Sequencing and library preparation of SLAM-seq was purchased from Single Cell Genomics Core, University of Eastern Finland. Sequencing service of ATAC-seq was purchased from BGI. Finally, we acknowledge the Turku Proteomics Facility’s service for mass spectrometry.

## Author contributions

JL led, managed and administered this study, proposed hypotheses, assessed the feasibility of chromatin accessibility and determined experimental methods, established and optimized ATAC-seq and metabolic labelling protocols, performed all experiments, collected all sequencing and mass-spectrometry samples, performed all bioinformatics analyses, generated and assembled all figures, and wrote the original draft of the manuscript. HMI formulated the initial concept of metabolic labeling-based enrichment of RNA Pol II and *CDK12*-inactivation-induced alterations in chromatin-accessibility, drafted the initial protocol for metabolic labeling-based enrichment of RNA Pol II and supervised the study. JL and HMI both conceptualized the study, obtained resources, planned the experiments, interpreted the results, and finalized the manuscript.

## Ethics declarations

### Ethics approval and consent to participate

Not applicable.

## Consent for publication

Not applicable.

## Abbreviations

5-EU: 5-ethynyl uridine
AR: Androgen receptor
ATAC-seq: Assay for transposase accessible chromatin with high-throughput sequencing
ATM: Ataxia-telangiectasia mutated
BRCA1: Breast cancer type 1
BRD4: Bromodomain containing 4
CRPC: Castration-resistant PC
CDK: Cyclin-dependent kinase
CDK12: Cyclin-dependent kinase 12
CTD: C-terminal domain
CTCF: CCCTC-binding factor
DDB1: Damage-specific DNA binding protein 1
DDR: DNA damage response
DARs: Differentially accessible regions
DEGs: Differentially expressed genes
DTT: dithiothreitol
ELOA: Elongin A
FBS: fetal bovine serum
GO: Gene ontology
HR: Homologous recombination
IP: Immunoprecipitation
IPA: Intronic polyadenylation
iPOND: Isolation of Proteins on Nascent DNA
KEGG: Kyoto Encyclopedia of Genes and Genomes
KLF: Krüppel-like factor
MF: Molecular function
MS: Mass spectrometry
mCRPC: Metastatic castration-resistant prostate cancer
NHEJ: Mon-homologous end joining
PBS: Phosphate-buffered saline
PC: Prostate cancer
PRKDC: Protein kinase, DNA-activated, catalytic subunit
Ser2: Phosphorylating Serine-2
Ser5: Phosphorylating Serine-5
RICK: RNA interactome
RNA: Pol II RNA polymerase II
RAD51: DNA repair protein RAD51 homolog 1
TDP: Tandem duplicator phenotype
pTEFB: The positive transcription elongation factor
SLAM-seq: Thiol(SH)-linked alkylation for the metabolic sequencing of RNA
SLAM-DUNK: Thiol(SH)-linked alkylation for the metabolic sequencing of RNA-Digital Unmasking of Nucleotide conversions in K-mers
TF: Transcription factor
TSS: Transcription start sites
TRCs: Transcription-replication conflicts

## Supplementary figures and supplementary figure legends

**Supplementary figure 1.**
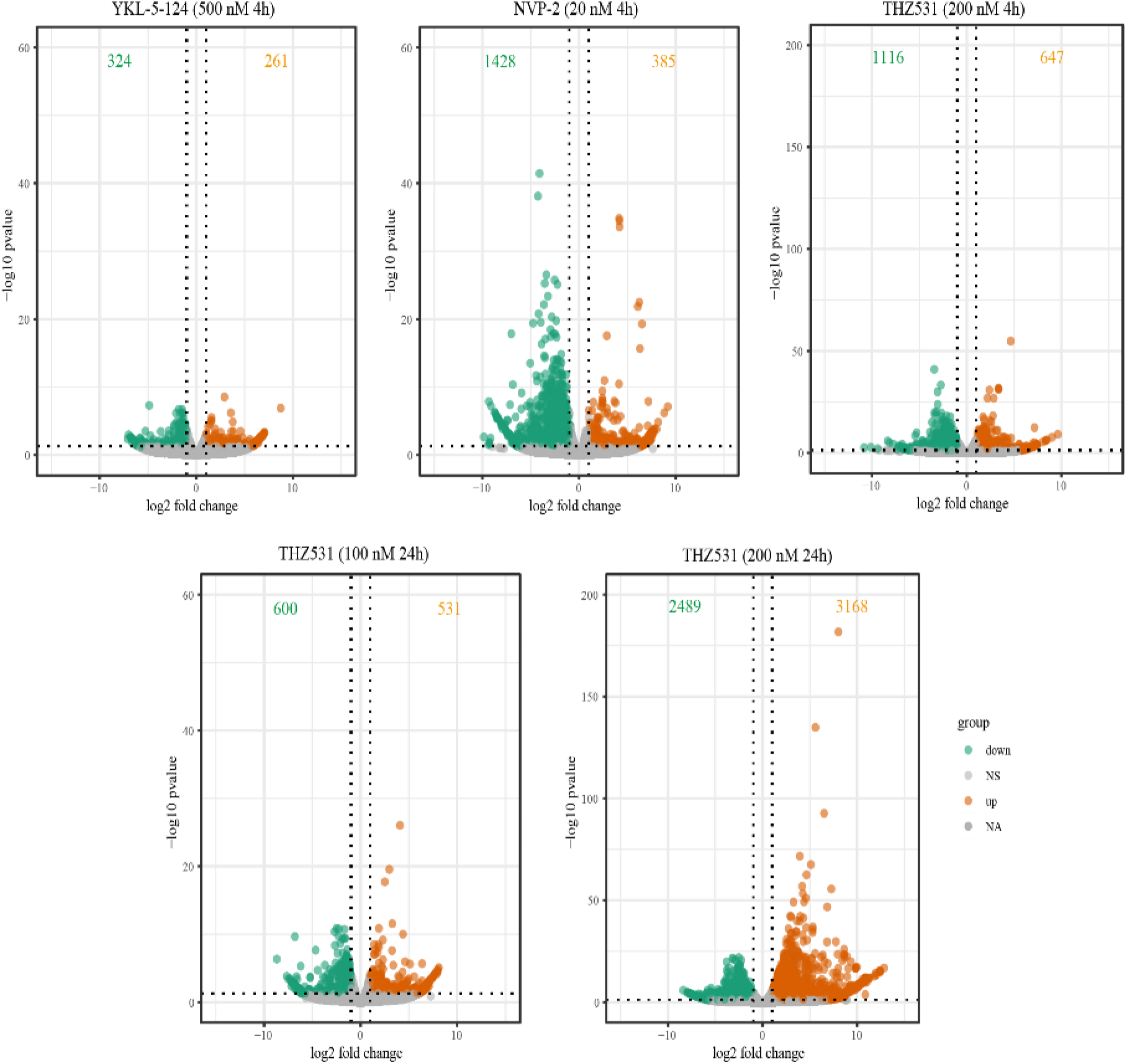
Differential gene expression analysis. Volcano plots showing differentially expressed genes (DEGs) in C4-2 cells treated with THZ531 (200 nM) for 4 h and 24 h. Volcano plots show significantly differentially expressed genes (DEGs) in C4-2 cells under different inhibitor treatments and at different treatment durations. Conditions: 500 nM YKL-5-124 (4 h), 20 nM NVP-2 (4 h), 200 nM THZ531 (4 h), 100 nM THZ531 (24 h), 200 nM THZ531 (24 h).

**Supplementary figure 2.**
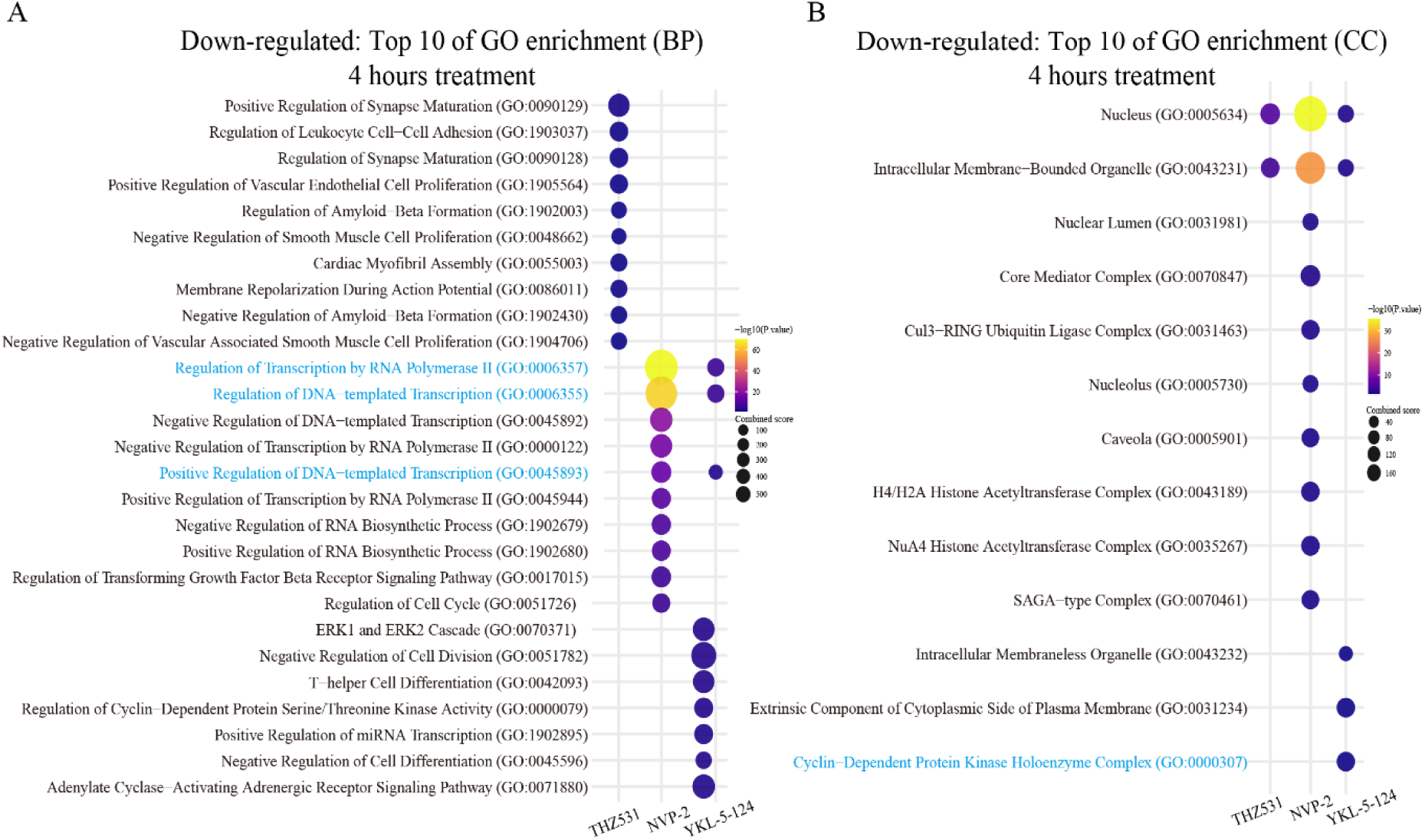
GO enrichment analysis was performed on downregulated genes after 4 hours of treatment with CDK7 (500 nM YKL-5-124), CDK9 (20 nM NVP-2), and CDK12 (200 nM THZ531) inhibitors for BP and CC pathways. **A** The dotplots show that in the BP pathway, downregulated genes were mainly enriched in after CDK7 and CDK9 inhibition. **B** The dotplot shows that in the CC pathway, the “Cyclin-Dependent Protein Kinase Holoenzyme Complex” was significantly enriched in downregulated genes under CDK7 inhibition. Pathways of particular interest are highlighted with blue color. GO: Gene Ontology, BP: Biological Process, CC: Cellular Component.

**Supplementary figure 3.**
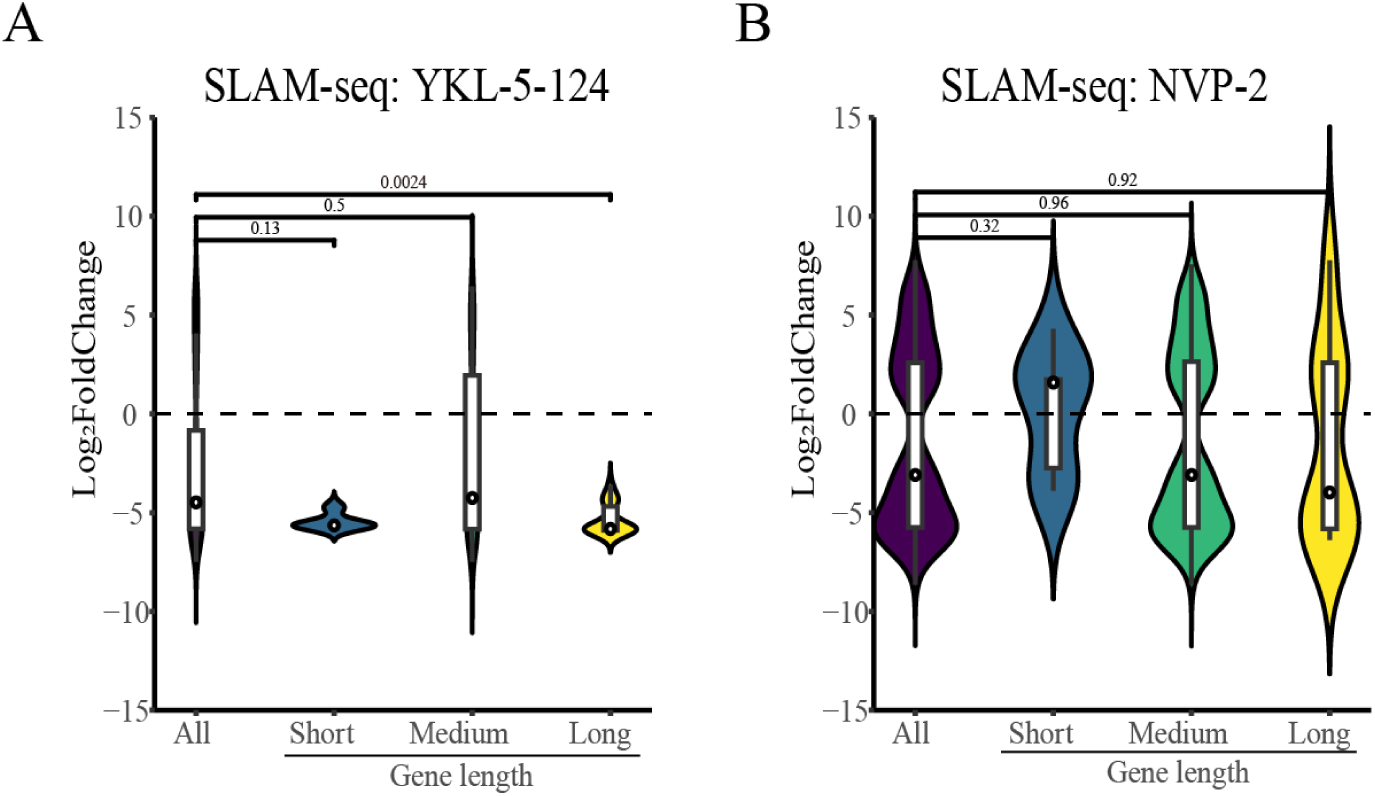
Length-dependent gene expression suppression by CDK7 and CDK9. **A** Violin plot results show that CDK7 suppression significantly downregulates the expression of long genes. **B** CDK9 inhibition does not significantly suppress the expression of long genes. Gene length stratification was consistent with our previous study [18], and significance was assessed using Student’s *t*-test. Gene length: short (<1 kb), medium (1–15 kb), and long (>15 kb).

**Supplementary figure 4.**
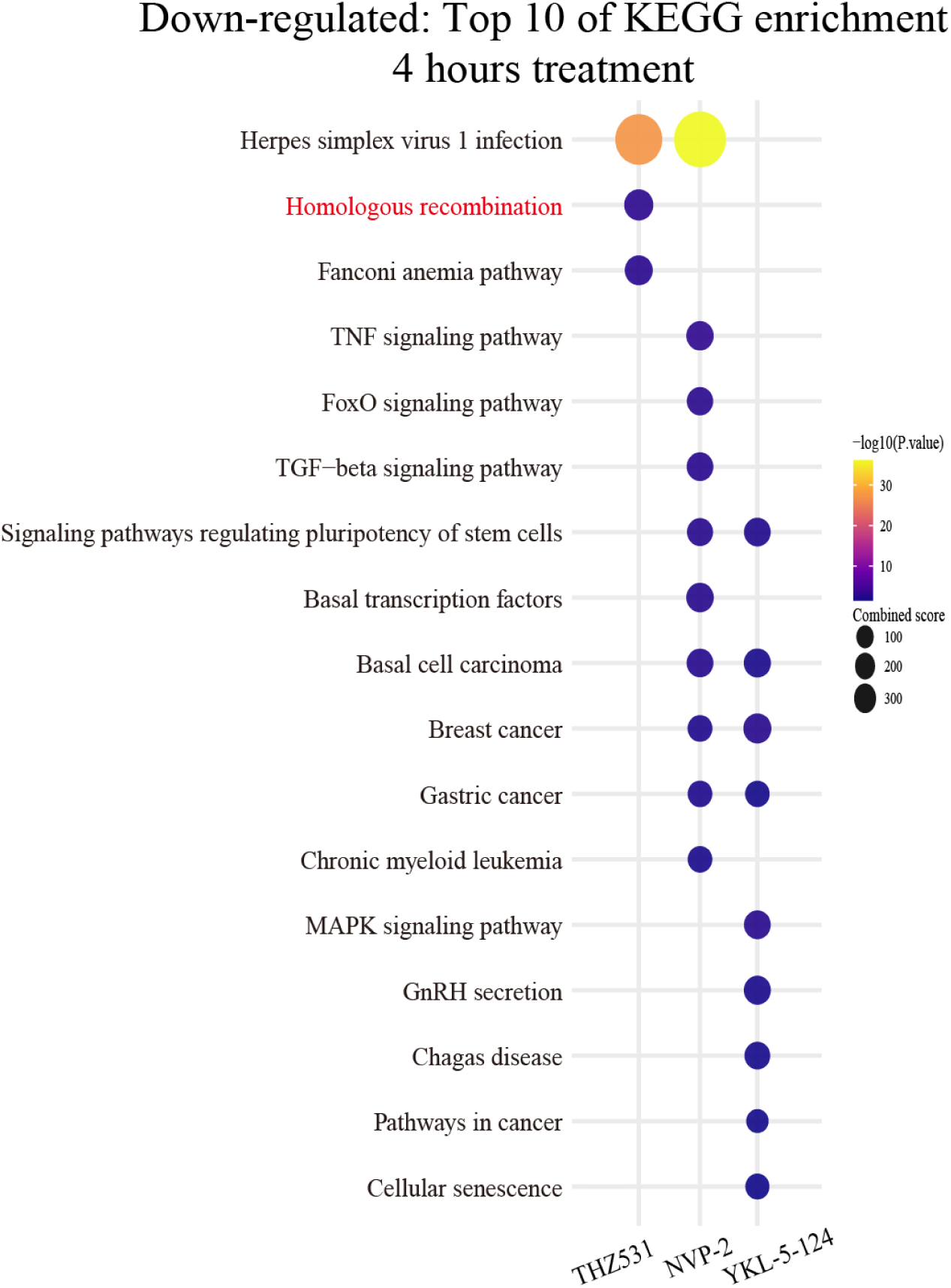
Genes significantly downregulated by CDK12/13 inhibition showed a significant enrichment of the homologous recombination pathway in KEGG analysis. The dotplot shows the enrichment analysis of KEGG for significantly downregulated genes after treatment with CDK7 (500 nM YKL-5-124), CDK9 (20 nM NVP-2), and CDK12/13 (200 nM THZ531) inhibitors for 4 hours. CDK12/13 inhibition significantly enriched the homologous recombination pathway, while CDK7 and CDK9 inhibition do not enrich.

**Supplementary figure 5.**
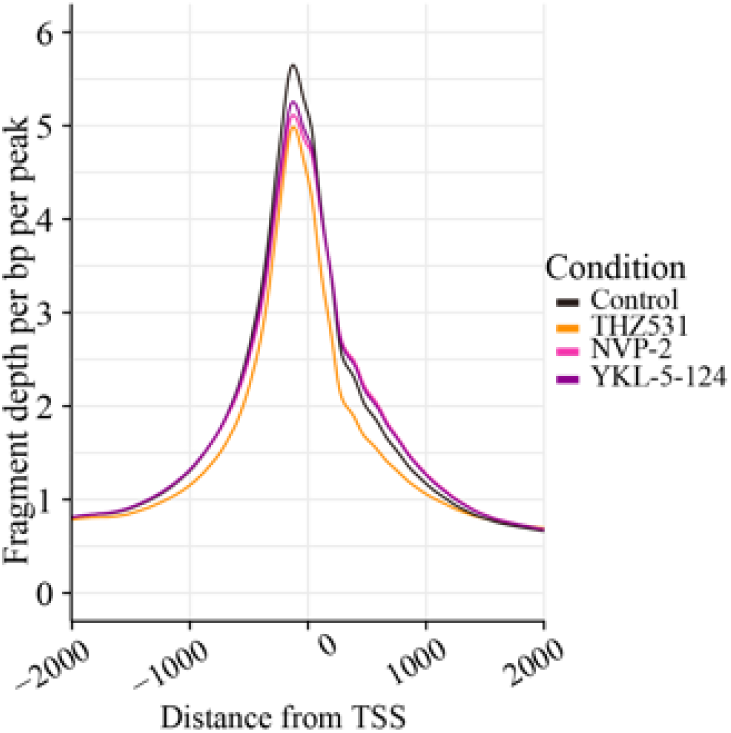
Comparative analysis of chromatin accessibility. The metagene plot compared changes in chromatin accessibility in the TSS region 4 hours after CDK7 (500 nM YKL-5-124), CDK9 (20 nM NVP-2), and CDK12 (200 nM THZ531) inhibition. The plot shows that inhibition of all transcriptional kinases results in chromosome closure. The chromosome openness ranking was Control > CDK7i > CDK9i > CDK12i.

**Supplementary figure 6.**
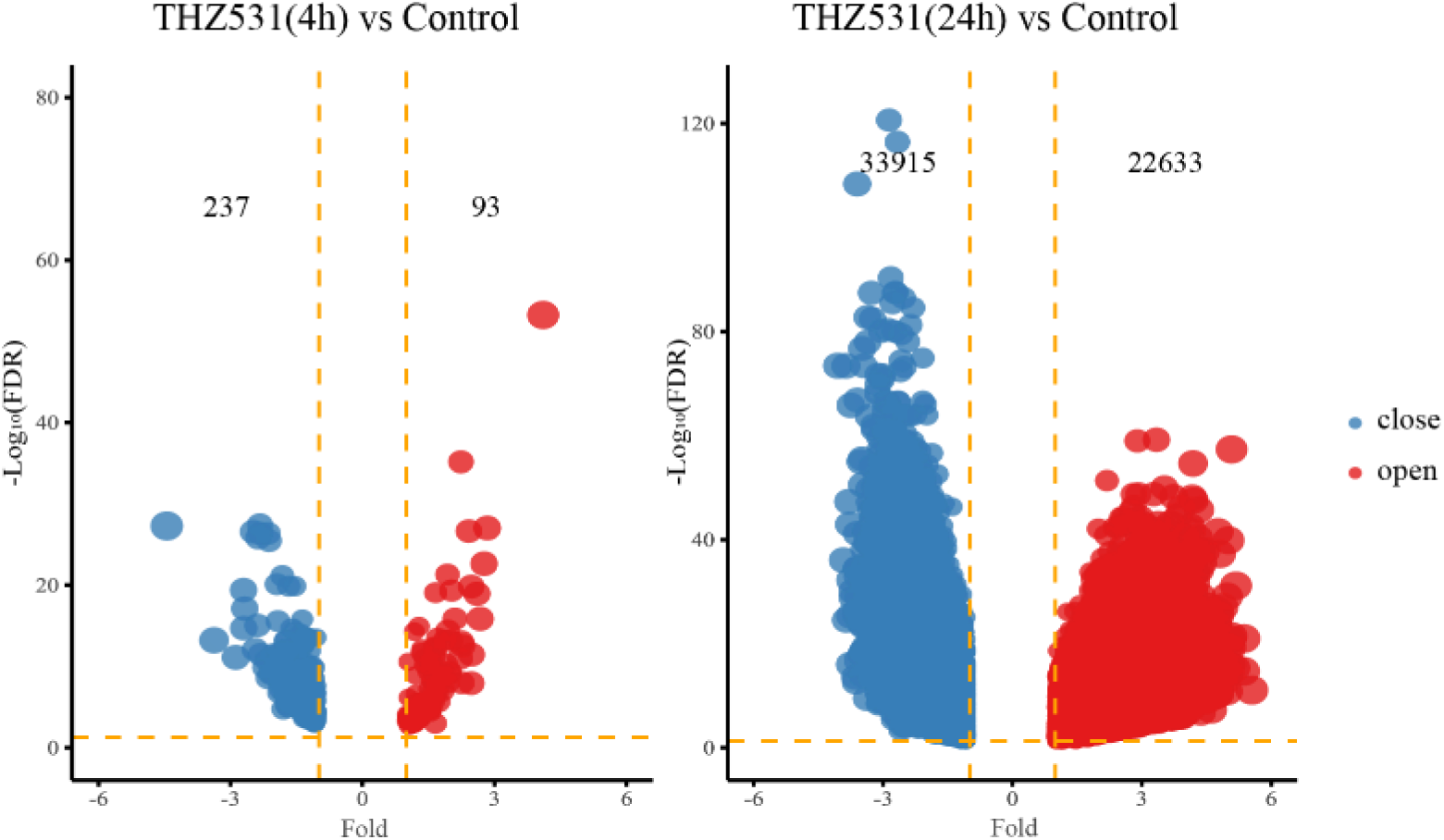
Analysis of differential accessibility under transcriptional kinase inhibition (CDK12/13). Volcano plots show significantly open and closed chromatin regions after 4 h and 24 h of treatment with 200 nM THZ531.

**Supplementary figure 7.**
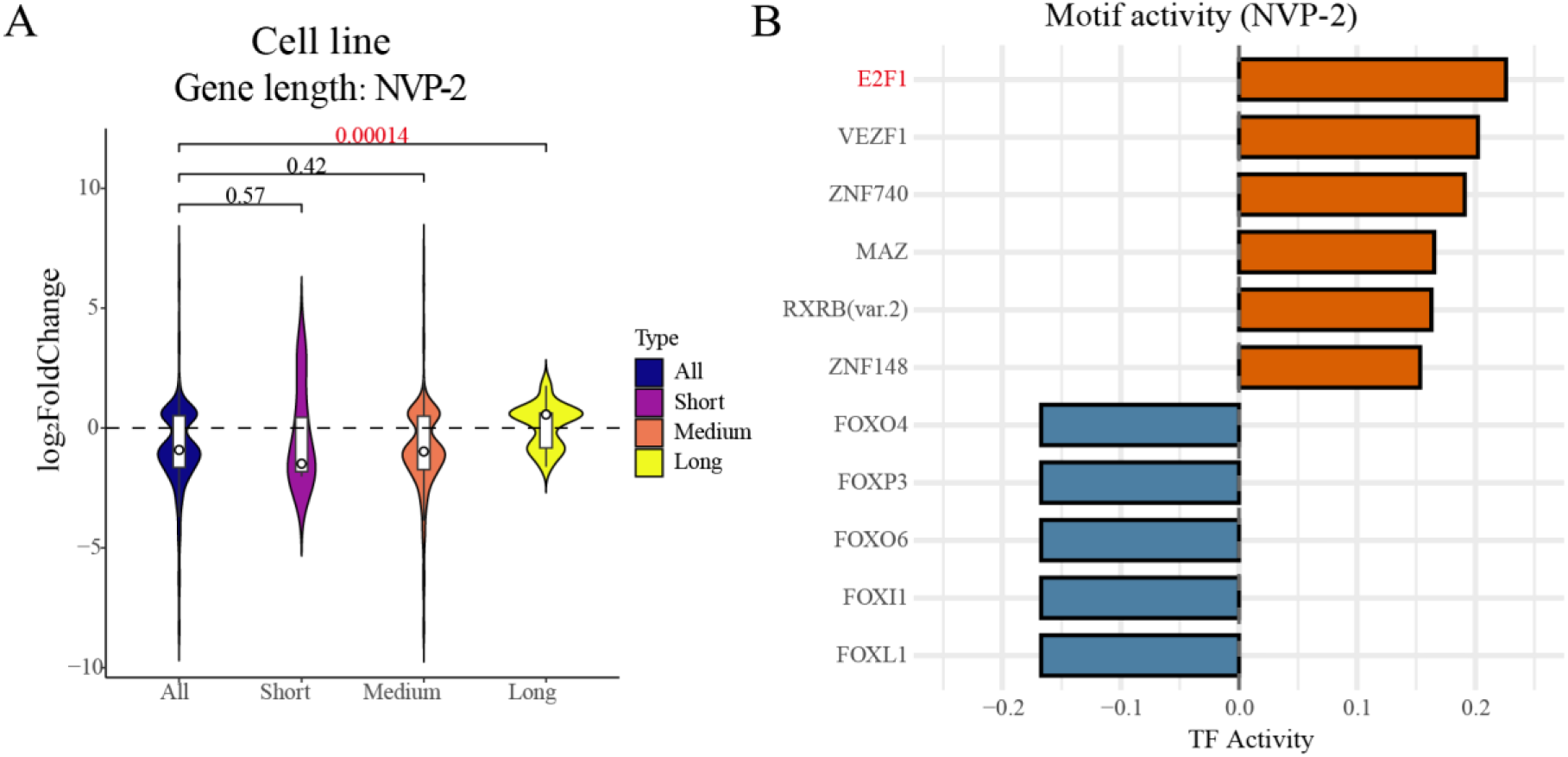
CDK9 repression increases chromatin accessibility of long genes. **A** Gene length analysis of differentially accessible regions with CDK9 inhibition (20 nM NVP-2 for 4 hours). The violin plot shows that CDK9 inhibition significantly upregulated the expression of long genes in differentially accessible genes. Significance was assessed using Student’s *t*-test. **B** Motif analysis of differentially accessible regions under CDK9 repression. The result shows significant enrichment of E2F transcription factors in NVP-2-induced open regions.

**Supplementary figure 8.**
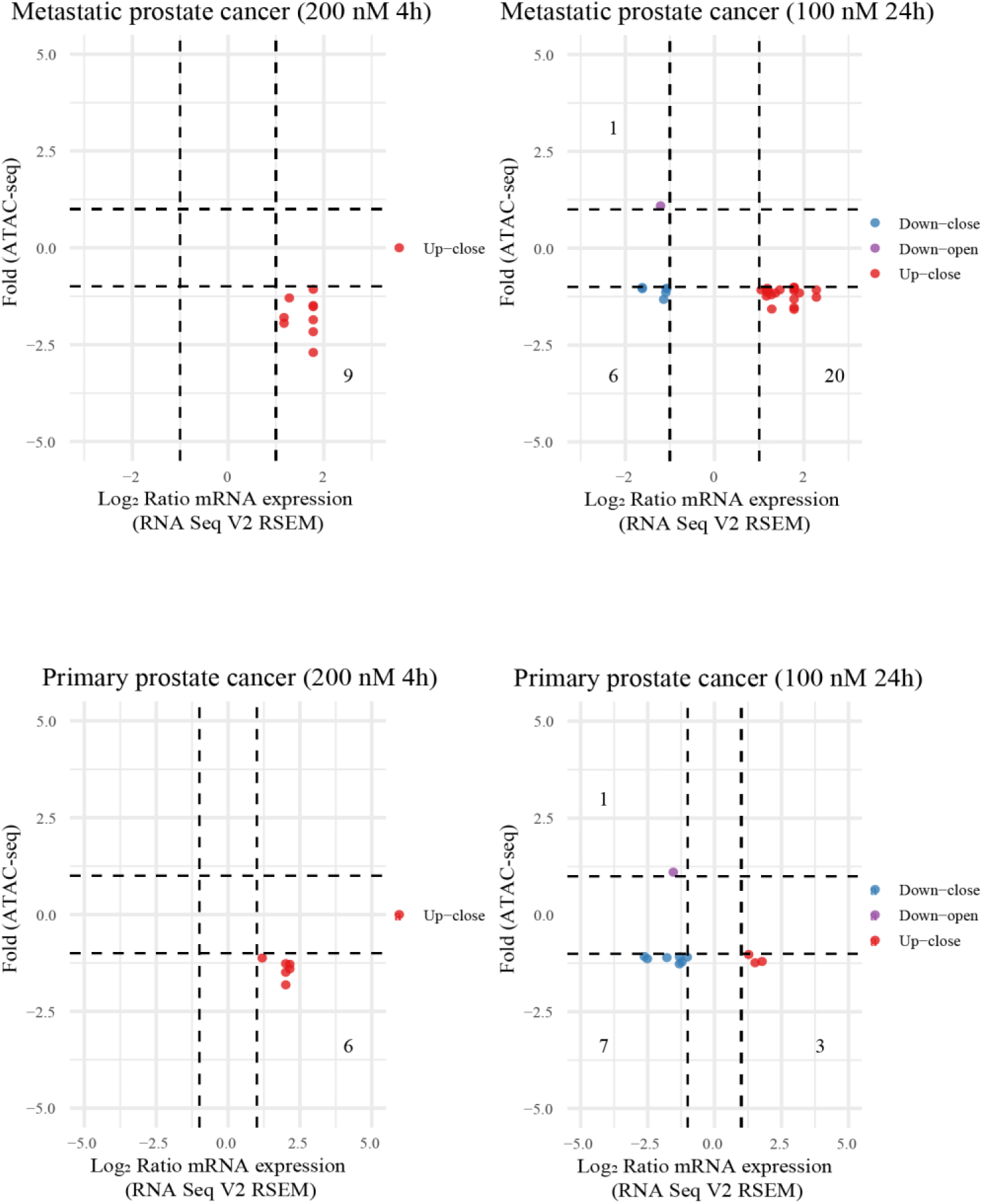
Integrative analysis of *in vitro* ATAC-seq data (CDK12/13 inhibition) with expression data from metastatic and primary prostate patient tumors from the TCGA database. The four-quadrant plot shows the data distribution of differentially accessible genes (4h: 200 nM THZ531 and 24h: 200 nM THZ531) with the expression profiles of TCGA data indicating patients with *CDK12* truncating mutation.

**Supplementary figure 9.**
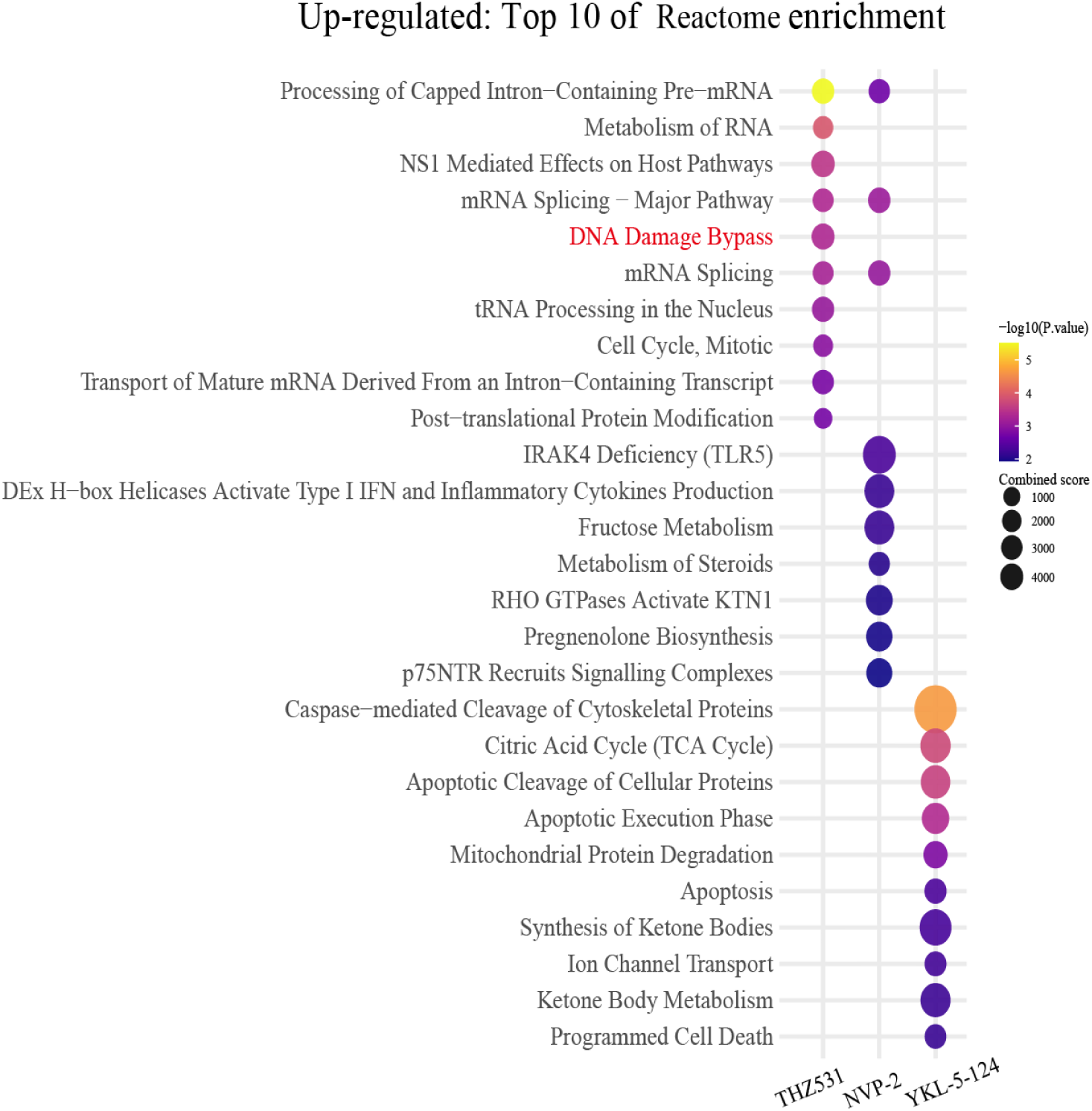
Reactome pathway enrichment analysis of genes significantly upregulated after treatment with CDK7, CDK9, and CDK12. This dotplot shows the reactome pathway enrichment analysis of genes significantly upregulated after 4 hours of treatment with CDK7 (500 nM YKL-5-124), CDK9 (20 nM NVP-2), and CDK12/13 (200 nM THZ531) inhibitors. Genes upregulated by CDK12/13 inhibition shows significant enrichment of the DNA damage bypass pathway, while CDK7 and CDK9 inhibition do not enrich for the same category.

**Supplementary figure 10.**
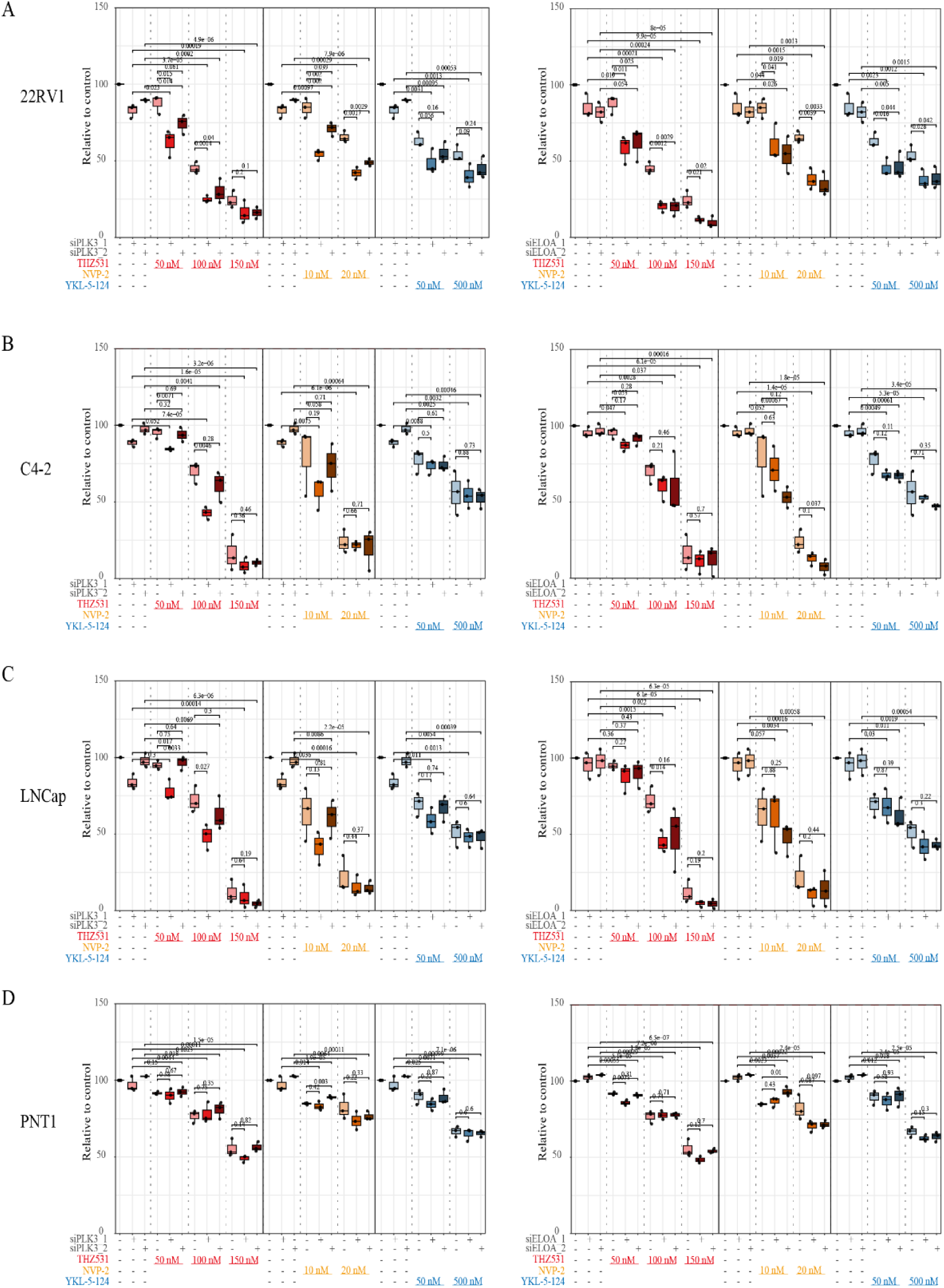
Functional validation of recruitment factors ELOA and PLK3. Cell viability was assessed using CellTiter-Glo 4 days after *ELOA* and *PLK3* knockdown, followed by inhibition with CDK7, CDK9, and CDK12 inhibitors for the last 3 days (the doses are indicated in the figure). Data were obtained from 3 biological replicates (each biological replicate contained 3 technical replicates), and the standard error of the mean was calculated. Significance was assessed using Student’s *t*-test. **A** Viability assay shows that knockdown of ELOA or PLK3 induce combinatorial anti-proliferative effects with CDK12, CDK9, and CDK7 inhibition, respectively, in the 22RV1 cell line. **B** and **C** Viability assays show that knockdown of ELOA or PLK3 do not induce combined lethality with CDK12, CDK9, and CDK7 inhibition in the C4-2 and LNCaP cell lines. **D** Viability assay shows that knockdown of ELOA or PLK3 do not trigger combined lethality with CDK12, CDK9, and CDK7 inhibition in the normal prostate cell line PNT1.

**Supplementary figure 11.**
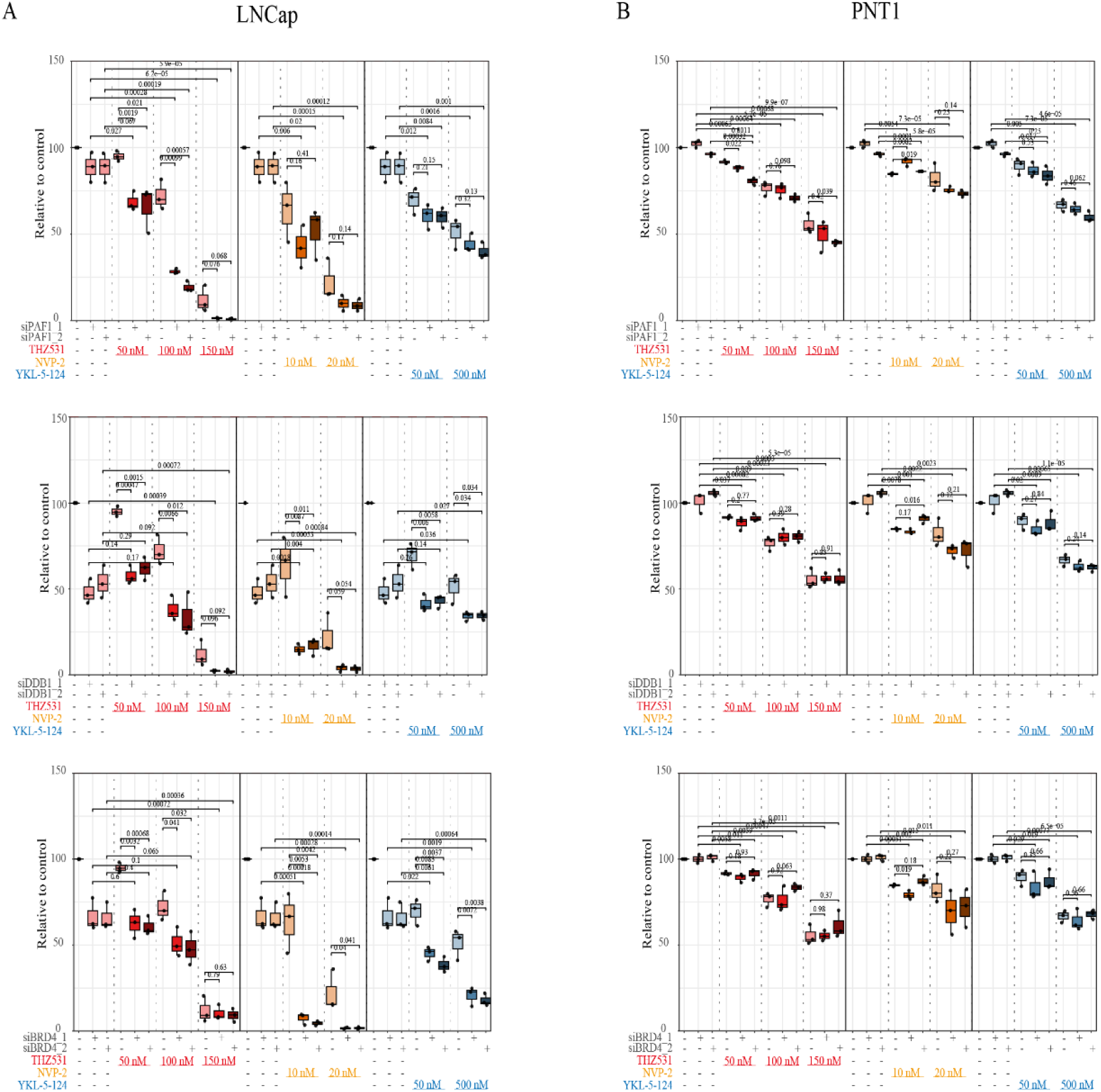
Functional validation and evaluation of recruitment factors DDB1, BRD4, and PAF1 in the prostate cancer cell line LNCaP and the normal prostate epithelia cell line PNT1. **A** Knockdown of PAF1 in androgen-dependent cell line LNCaP induces combined reduction in cell viability with CDK12 inhibition, but does not trigger this combined effect with CDK9 and CDK7 inhibition. Knockdown of DDB1 in LNCaP induces combinatorial anti-proliferative effects with CDK12/13, CDK9, and CDK7 inhibition. Knockdown of BRD4 in LNCaP decreased cell viability with CDK9 and CDK7 inhibition but this effect is not significant with 150 nM treatment (CDK12/13) inhibition. **B** Knockdown of DDB1, BRD4, and PAF1 in the cell line immortalized from normal prostate epithelia, PNT1 does not induce significant combinatorial anti-proliferative effects with CDK12/13, CDK9, and CDK7 inhibition. Data were obtained from 3 biological replicates (each biological replicate contained 3 technical replicates), and the standard error of the mean was calculated. Significance was assessed using Student’s *t*-test.

---

**Supplementary Table 1.**
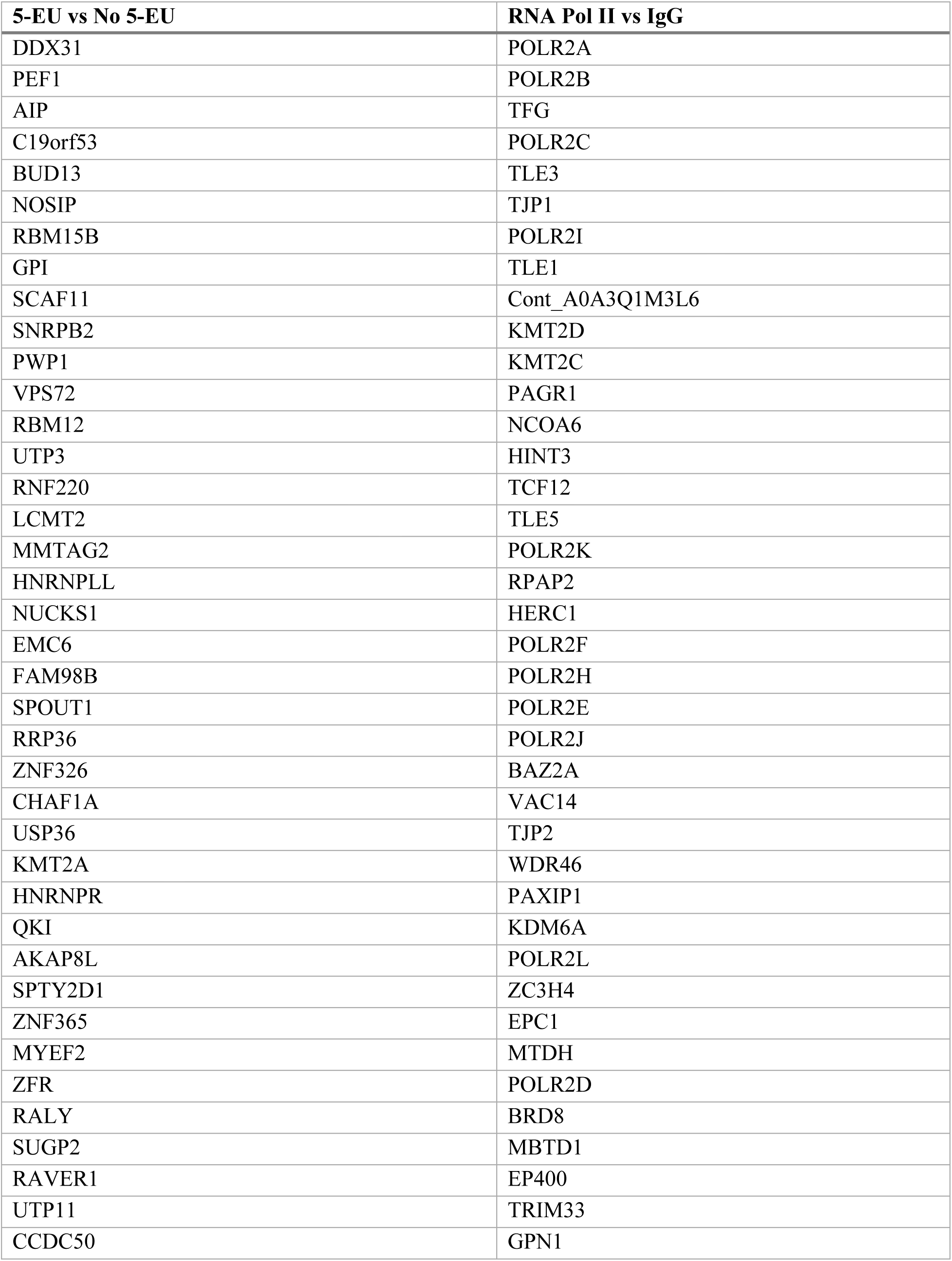
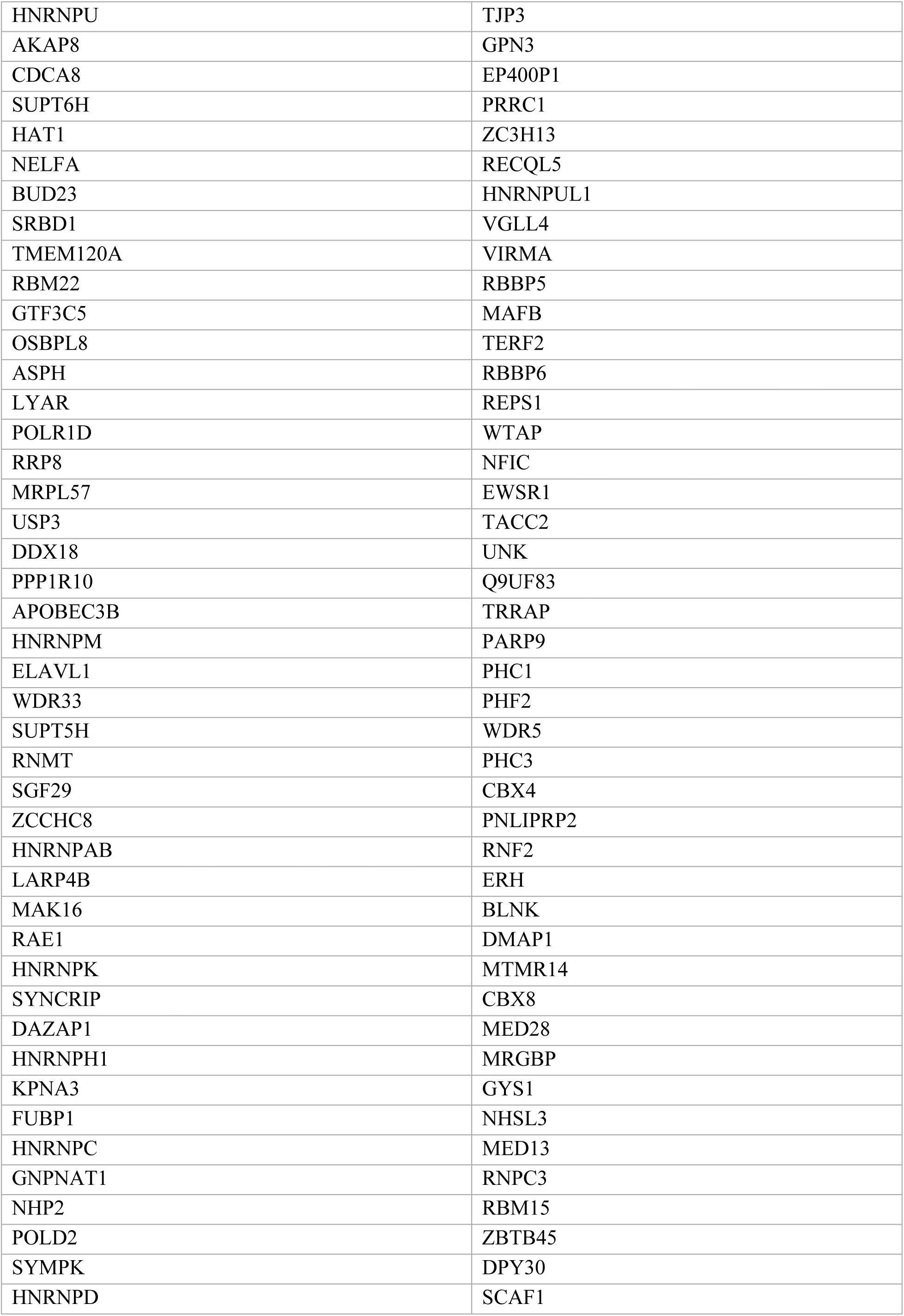
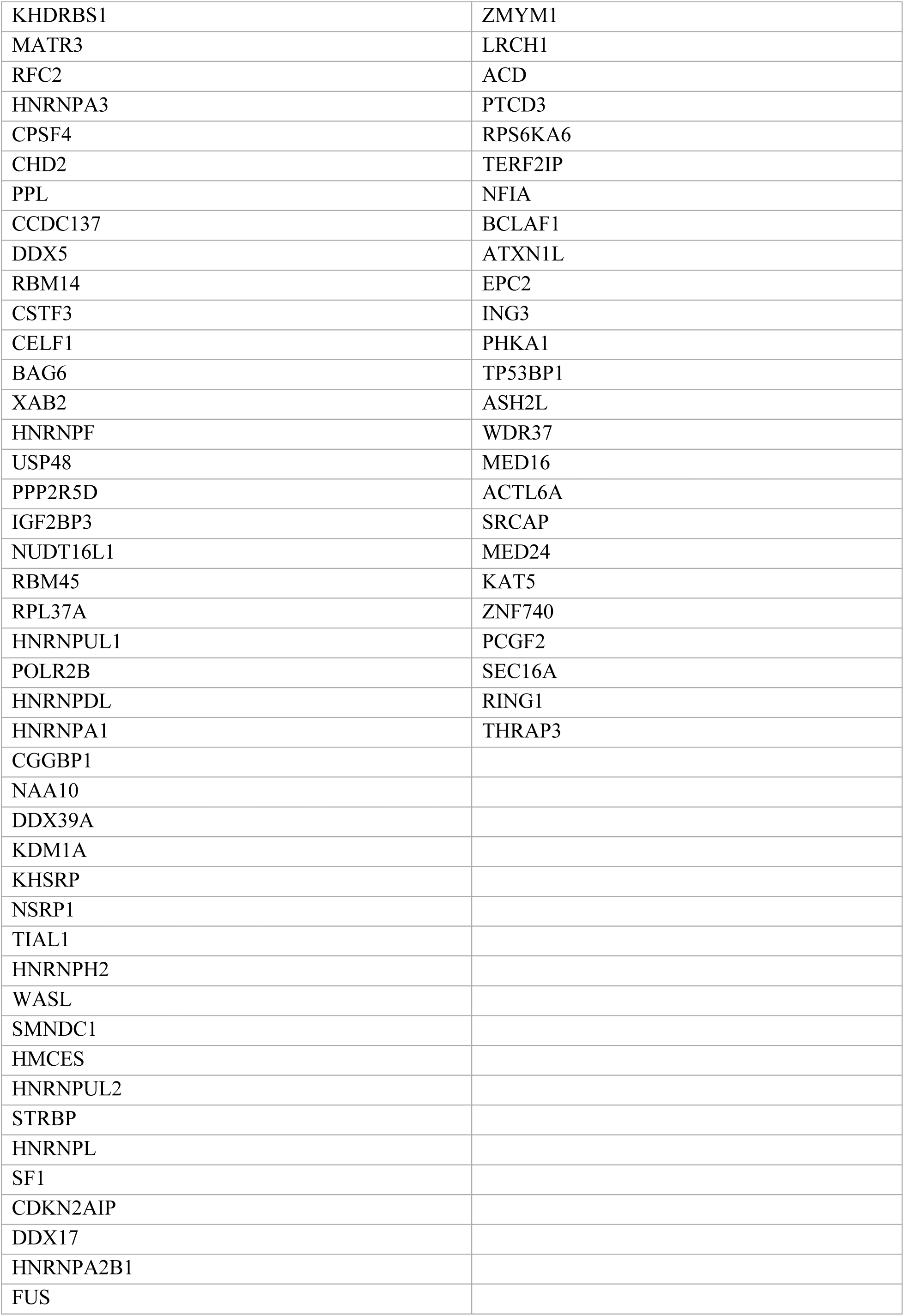
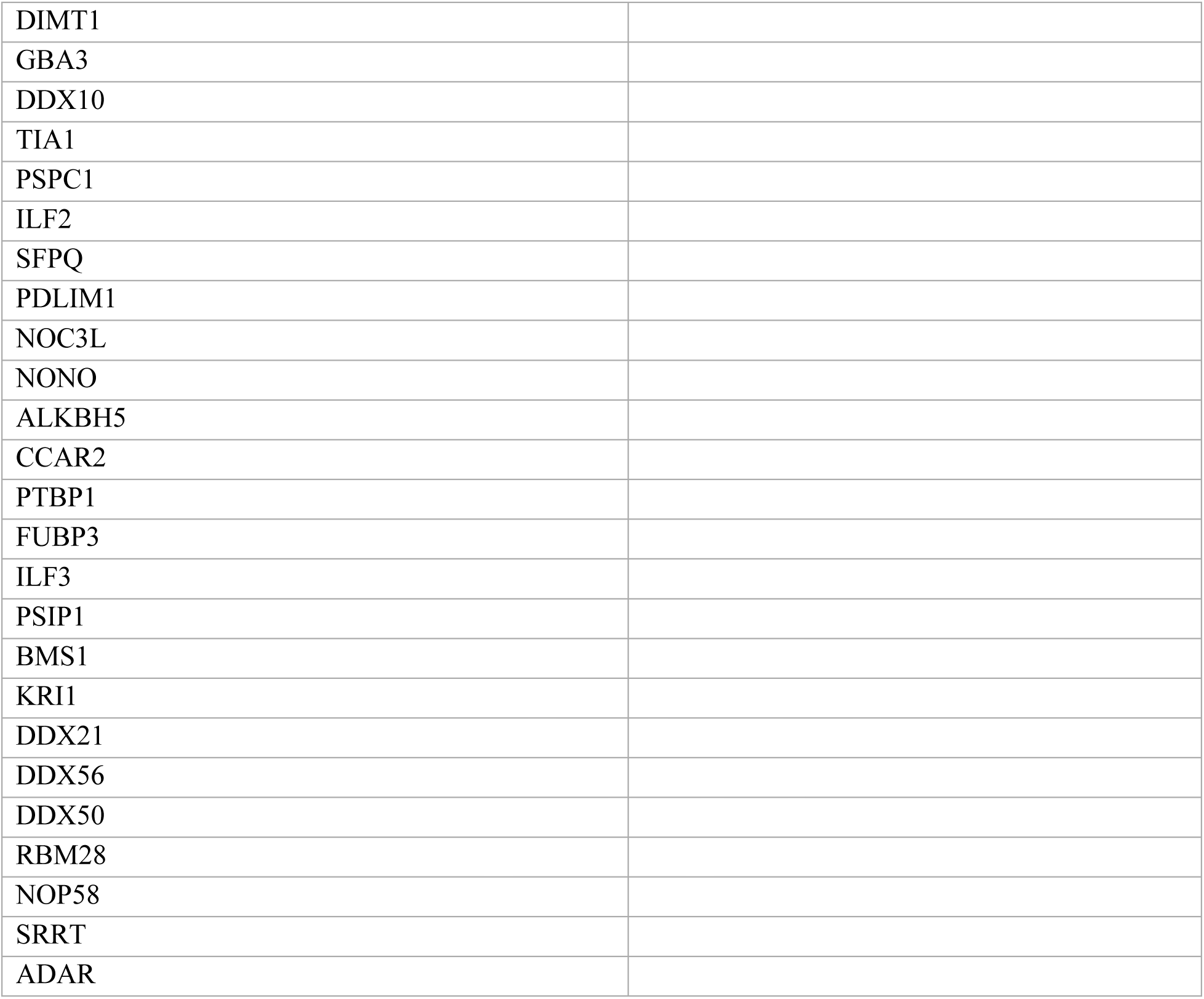
This table shows proteins that are significantly enriched by 5-EU labelling (5-EU vs No 5-EU) and proteins that are significantly enriched by RNA polymerase II immunoprecipitation (RNA Pol II vs IgG).

